# Protein Sequence Modelling with Bayesian Flow Networks

**DOI:** 10.1101/2024.09.24.614734

**Authors:** Timothy Atkinson, Thomas D. Barrett, Scott Cameron, Bora Guloglu, Matthew Greenig, Louis Robinson, Alex Graves, Liviu Copoiu, Alexandre Laterre

## Abstract

Exploring the vast and largely uncharted territory of amino acid sequences is crucial for understanding complex protein functions and the engineering of novel therapeutic proteins. Whilst generative machine learning has advanced protein sequence modelling, no existing approach is proficient for both unconditional and conditional generation. In this work, we propose that Bayesian Flow Networks (BFNs), a recently introduced framework for generative modelling, can address these challenges. We present ProtBFN, a 650M parameter model trained on protein sequences curated from UniProtKB, which generates natural-like, diverse, structurally coherent, and novel protein sequences, significantly outperforming leading autoregressive and discrete diffusion models. Further, we fine-tune ProtBFN on heavy chains from the Observed Antibody Space (OAS) to obtain an antibody-specific model, AbBFN, which we use to evaluate zero-shot conditional generation capabilities. AbBFN is found to be competitive with, or better than, antibody-specific BERT-style models, when applied to predicting individual framework or complimentary determining regions (CDR).

## Introduction

Proteins drive nearly every process in biological systems, playing a central role in the operations of both healthy and pathological functions. However, despite this only a small fraction of the possible proteome - characterised by the vast combinatorial complexity of possible amino acid sequences - has been explored with vast regions of the theoretical space not yet having been observed in natural proteins [7, 46]. This represents not only a gap in our fundamental knowledge of biological processes, but also potentially untapped opportunities for developing novel functional and therapeutic proteins [10, 17].

In recent years, machine learning (ML) has emerged as a powerful tool to bridge this gap, becoming an integral part of computational biology. Techniques initially developed for natural language processing (NLP) have proven particularly impactful. Drawing on parallels between modelling sequences of words and amino acids (the ‘language of proteins’), these self-supervised models learn to generate novel sequences from corpora of unlabelled data. For example, BERT-style models – such as the Evolutionary Scale Modeling (ESM) series [35, 36, 49] – are trained to predict masked amino acids based on the context of the rest of the sequence. However, whilst being trained to conditionally generate missing parts of a sequence; BERT models are ill-suited for the generation of entirely new protein sequences. Instead, they are primarily used to generate embeddings used for various downstream tasks such as folding [35], inverse folding [28], contact prediction [48] and various other applications [8, 40, 50].

As generative pretrained transformer (GPT) models have emerged as the dominant paradigm in NLP [20, 61], equivalent autoregressive protein language models – such as ProtGPT2 [19], ProtGen [41] and RITA [25] – have been explored for *de novo* protein sequence generation. These autoregressive models build sequences from left to right with each amino acid conditioned only on the preceding partial sequence. However, as the function of proteins is determined by their 3D structure, regions of interest are rarely contiguous segments at a terminus of the chain. Therefore GPT models are unsuitable for many practical design tasks. Indeed, this bi-directional co-dependency of functional proteins, between different positions in the sequence, makes it far from clear that an autoregressive prior is an optimal choice for protein generation.

Given the limitations of treating protein sequences strictly as linguistic constructs, recent efforts have explored generative techniques from other domains. Notably, diffusion models [27, 31], which excel in image generation [13], have shown promise in structural protein modeling through adaptations like RFDiffusion [64] and AlphaFold3 [2]. However, diffusion models are tailored for continuous variables and attempts to adapt these techniques to handle the discrete nature of amino acid sequences (i.e. ‘discrete diffusion’) are still being refined. Indeed, no method has proved effective at both *de novo* unconditional generation and arbitrary conditional generation of novel protein sequences.

Here, we propose that recently developed Bayesian Flow Networks (BFNs) [23] can address the above mentioned challenges. BFNs are generative models that do not enforce a specific decomposition of the joint distribution over all variables. Moreover, rather than modelling the data directly, BFNs operate on the continuous parameters of a distribution over the data. As such, they naturally handle non-continuous data modalities, including discrete variables, as a first-class citizens. As they have only recently been proposed, BFNs have not been tested across the the same range of tasks as the aforementioned generative approaches. However, in this work we demonstrate that they can effectively handle the complexity of protein sequence modelling realising leading performance whilst unifying unconditional and conditional generation.

Specifically, we present ProtBFN, a 650 M parameter model trained for *de novo* generation of protein sequences. The generated proteins match the known natural distribution in terms of direct metrics such as amino acid propensity and sequence length and computed properties; whilst being globally coherent, globular sequences. Compared to state-of-the-art autoregessive and discrete diffusion models, we find ProtBFN’s samples are more natural whilst covering more of the known proteome. Moreover, ProtBFN is also capable of exploring unseen regions of the protein space; generating novel sequences with low identity but nevertheless high structural similarity to the natural motifs found in databases such as CATH [54].

Finally, we fine-tune ProtBFN to yield an antibody-specific model, AbBFN. Consistent with our findings for ProtBFN, AbBFN is able to unconditionally generate highly plausible heavy chain candidates. To validate the zero-shot conditional generation capabilities of our model, we then apply AbBFN to the task of inpainting individual CDR and framework regions of existing VH chains. Despite having only been trained for unconditional generation, AbBFN recovers the underlying amino acids following known patterns of variability and matches the performance of leading BERT-style transformer models trained specifically for this task. Moreover, the flexible conditional generation of BFNs allows AbBFN to retain strong amino acid recovery rates, and outperform the BERT models, when the masking region is increased to include all frameworks regions.

## Results

### A Bayesian Flow Network for Protein Sequences

Generative models aim to capture the joint distribution *p*(**x**) of a dataset, where each sample comprises *N* variables denoted as **x** = [*x*_1_, …, *x*_*N*_ ]. Typically, these models are trained to reconstruct the original data from noised or corrupted samples. Traditional sequence modeling techniques, such as GPT and BERT, manipulate subsets of these variables to create noised training targets. Specifically, GPT models the distribution autoregressively 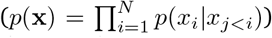 while BERT generates data conditionally based on non-masked variables

In contrast, BFNs uniformly apply noise across all variables, similar to variational diffusion models. The generation of a new sample unfolds as a continuous-time denoising process, initiating from a random prior and concluding with a well-defined sample. Unlike traditional diffusion techniques that directly model the *data*, **x**, BFNs adjust the parameters of the data’s *distribution, θ* = [*θ*_1_, …, *θ*_*N*_ ], where *p*(*x*_*i*_ | *θ*_*i*_) governs the distribution over the *i*^th^ variable. This distinction is crucial, especially for handling discrete data where variables are categorically correct or incorrect. This lack of intermediate gradations makes defining a smooth denoising process over the data (i.e. with the data become monotonically more accurate) a significant challenge in the application of traditional diffusion methods.

### Training

The denoising process used to generate samples can be viewed as a communication protocol (see Fig. 1), where ‘Alice’ describes a ground-truth data point (e.g. a sequence of amino acids) to ‘Bob’ by sending a series of noisy observations, **y**^(1:*i*)^ ≡{**y**^(1)^, … **y**^(*i*)^ }, that become increasingly more informative. Training a BFN is then equivalent to Bob learning a model that, given a series observations with known noise, predicts a data distribution that best matches next observations, **y**^(*i*+1)^. Concretely, this training process at step *i* applied to protein sequences is as follows.

1. Alice sends a noisy observation (**y**^(*i*)^) of a protein (**x**) from the *sender distribution* 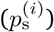 to Bob.
2. Bob summarises all noisy observations, **y**^(1:*i*)^, on a per-amino-acid basis via Bayesian inference, to obtain the parameters of an *input distribution θ*^(*i*)^.
3. These parameters are input into a neural network Φ, which predicts an *output distribution* 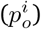. This step models the joint distribution of amino acids, refining the single-variable Bayesian inferences.
4. To obtain Bob’s belief over the form of the next noisy observation, (**y**^(*i*+1)^), the output distribution is noised in an equivalent manner to the ground truth data, to obtain the *receiver distribution* 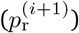.

**Figure 1.**
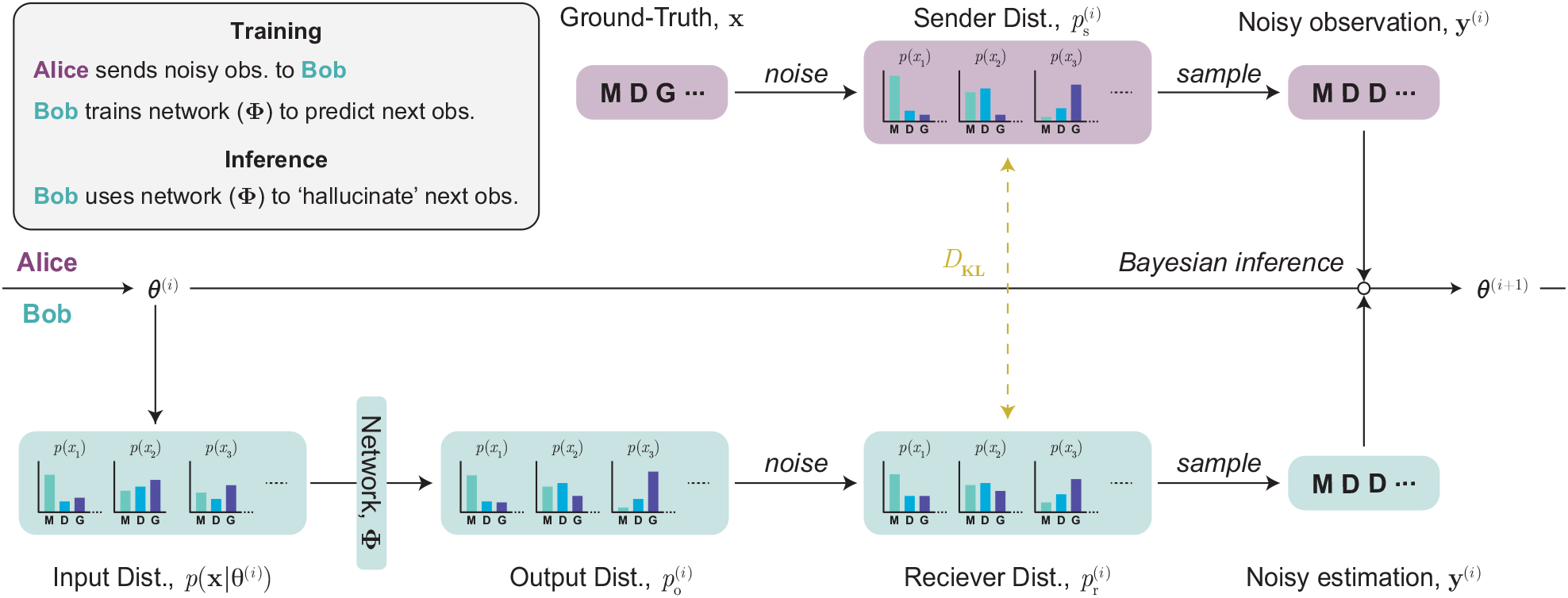
Application of a Bayesian Flow Network (BFN) to protein sequence modelling. BFN’s update parameters of data distribution, *θ*, using Bayesian inference given a noised observation, **y** of a data sample. When applied to protein sequence modelling, the distribution over the data is given by separate categorical distributions over the possible tokens (all amino acids and special tokens such as <pad>, <bos>, and <eos>) at each sequence index. During training, ‘Alice’ knows a ground truth data point **x**, and so *θ* can be directly updated using noised observation of **x**. ‘Bob’ trains a neural network to predict the ‘sender’ distribution from which Alice is sampling these observations at each step (i.e. to predict the noised ground truth). During inference, when Alice is not present, Bob replaces noised observations of the ground truth with samples from the ‘reciever’ distribution predicted by the network.

Minimising the difference between the predicted and true distribution over the next data samples in an *N* step generation process is then given by the training objective;

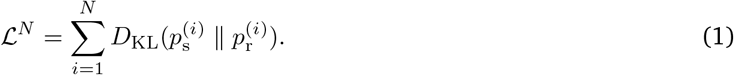

It is possible to derive a continuous-time loss function ℒ^*∞*^ in the limit of *N* → ∞ [23] which is used in practice (see Methods for details).

### Inference

When sampling unconditionally full protein sequences from the model, there is no fixed ground-truth. Instead, Bob effectively ‘hallucinates’ Alice with the previous step’s receiver distribution takes on the role of the sender distribution 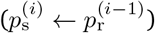. Repeating this process *N* times generates samples from the learned distribution.

Whilst a naive approach for conditional generation is to use the sender distribution at variables with a ground truth and the receiver distribution elsewhere, we found that this does not converge to the true conditional distribution. Instead, we combine this approach with sequential Monte Carlo (SMC) sampling; which ensures that the sampled variables are consistent with fixed variables under the learned joint distribution. Full details on the sampling methodology are found in the Methods.

### ProtBFN: A foundational generative model for protein sequences

Using the aforementioned methodology, we trained a 650 M parameter model, ProtBFN, on a curated dataset of sequences that span the known proteome. This dataset considers non-hypothetical protein sequences UniProtKB [1] and we refer to this curated and clustered dataset as UniProtCC. In assessing the capabilities of ProtBFN, we compare against leading autoregressive and discrete diffusion models for protein sequence generation – ProtGPT2 [19] and EvoDiff [3], respectively. We note that both ProtGPT2 and EvoDiff are trained on the UniRef50 dataset which, unlike UniProtCC, includes proteins of hypothetical or unknown existence. By ensuring ProtBFN trains on only sequences with higher confidence in their existence we ensure it remains focused on the biologically relevant manifold of possible proteins. Our evaluation analyses 10 000 protein sequences generated by each model and equivalenltly sized subsets randomly extracted from both UniProtCC and UniRef50 datasets. Extended details of the data curation, training procedure and sampling methodology are provided in the Methods.

### Learning to generate natural protein sequences

A primary goal of ProtBFN is to learn the distribution of natural proteins. Although no single metric can definitively assess the ‘naturalness’ of generated protein samples, various statistical and biophysical properties can be computed and compared against the expected natural distributions. To this end, Fig. 2 presents a selection of such experiments from which we can infer that ProtBFN not only matches the natural distribution on which it was trained, but does so more faithfully than the autoregressive and discrete diffusion baselines on their respective training dataset.

**Figure 2.**
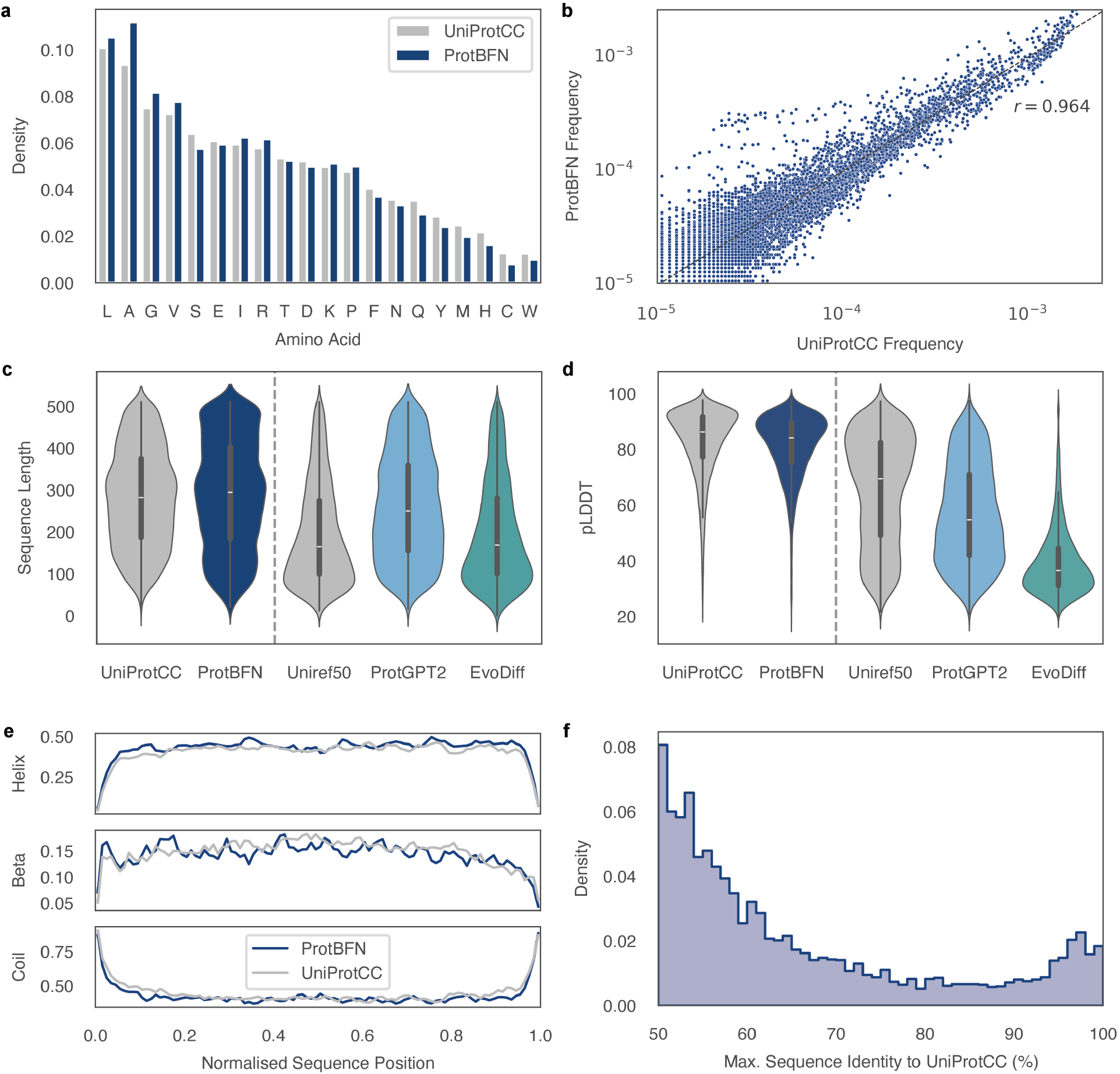
ProtBFN is an effective *de novo* generator of protein sequences; generating novel protein sequences that in distribution under both local and global metrics. The local metrics considered are amino acid (**a**) and (**b**) oligomer frequencies; both of which show a strong match to the training distribution of the model. The 50 256 oligomers used are taken from the vocabulary of ProtGPT2 [19] and the plot has overlaid a linear correlation fit and the associated Pearson correlation coefficient. Metric computed as functions of the entire protein sequence are used to assess the global coherence of the generations. Both the sequence lengths (**c**) and predicted mean local distance difference test (pLDDT) scores from ESMFold [36] (**d**) are shown. Predictions with pLDDT*>*70 are generally considered high confidence. Also included are a baseline autoregressive, ProtGPT2 [19], and discrete diffusion, EvoDiff [3], model. This structural analysis is extended to show the secondary structure along the length of the generated sequences as predicted by NetSurf P-3.0 [29] (**e**). In all cases, ProtBFN is seen to well match the natural training distribution UniProtCC. Additional results, including equivalent figures for baseline models and metrics not presented here can be found in Supplementary Section A. Finally, the maximum sequence identity of the ProtBFN generated sequences to the UniProtCC training data is plotted (**f**) to demonstrate that the training data is not being memorised.

A direct test of the plausibility of ProtBFN’s generated set of protein sequences is to examine the frequency with which amino acids and oligomers occur as these are strongly related to the robustness of the genetic code [21]. These measures (Fig. 2a and b) are well aligned with the UniProtCC distribution. However, whilst these frequency metrics are indicative they do not alone confirm the coherence of individual samples, therefore we also compute per-sample metrics. ProtBFN’s generated sequences have a natural-like distribution of lengths (Fig. 2c) which is another indicator of our ability to model the underlying distribution. In contrast, ProtGPT2 significantly diverges from the UniRef50 under this metric while EvoDiff artificially matches the target distribution because sequence length is pre-selected before generation. Additional results comparing biophysical properties of the generated sequences are detailed in Supplementary Section A.

For structural analysis we first leverage NetSurf P-3.0 [29] to annotate each residue in the sequences with structural information. We again find that ProtBFN exhibits equivalent annotations to the natural distribution, even when considering how these properties vary as a function of position in the sequence (Fig. 2d and Supplementary Section A). To then evaluate the overall structural properties, we use the mean predicted local distance difference test (pLDDT) of ESMFold [36] as a measure of confidence in the predicted structure (Fig. 2e). As interactions between residues far apart within in a protein sequence are critical in determining overall structure; higher pLDDT scores indicate that ESMFold can identify similar interactions within the generated samples as were seen in the training data. Indeed, we observe that ProtBFN consistently produces sequences which obtain high pLDDT scores, closely matching those of naturally occurring proteins. The sensitivity of this metric to the global coherence of a protein is evidenced by ProtGPT2 and EvoDiff’s divergence from their training distribution; even when accounting for the greater number of lower scored proteins in UniRef compared to UniProtCC.

Finally, to confirm ProtBFN is generating novel proteins rather than memorizing training data, we search for the nearest match for each generated sequence within the UniProtCC training data (Fig. 2f). Results show that generated sequences are highly likely to be novel, with 4444 (8851 and 9489) of the generated samples having sequence identity to the nearest match being less than 50 % (80 % and 95 %), respectively.

### ProtBFN effectively covers the known proteome

Broad coverage of the proteome ensures that a model has learned the diversity of protein sequences, and thus holds the potential for developing a wide array of functional proteins. Having confirmed the naturalness and novelty of the sequences generated by ProtBFN, we next assess the proteome coverage provided by these sequences. To do so, we first use `mmseqs2` [56] to align generated sequences with UniRef50 (Table 1), as UniRef50 represents a non-redundant set of protein clusters spanning the known protein diversity.

**Table 1.** ProtBFN generates more natural and diverse proteins than baseline meothds. 10 000 generated sequences from each model are aligned against UniRef50 with above 50 % sequence identity bet ‘ between a sequence and cluster counted as a ‘hit’. ProtBFN sequences are found to align with known clusters significantly more often than those from baseline models. The coverage score (see Methods for details) is proposed as a measure of the diversity of different clusters hit by the sequences. Higher scores corresponding to broader coverage of the target clusters.

| Model | Cluster hits | Coverage score |
| --- | --- | --- |
| ProtBFN | **69.7 %** | **0.544** |
| ProtGPT2 | 15.7 % | 0.095 |
| EvoDiff | 2.6 % | 0.034 |

69.7 % of ProtBFN’s of sequences are found to align (≥50 % sequence identity) with a known UniRef50 cluster, a significantly higher proportion than ProtGPT2 and EvoDiff despite that fact that these models are trained uniformly on the UniRef50 clusters. As the number of sampled sequences (10 000) is far smaller than the number of UniRef50 clusters (∼65M), we measure coverage as the ratio of observed to expected unique clusters hit if drawing 10 000 samples from a model’s training distribution. Calculation details of this proposed *coverage score* are provided in the Methods and empirically we find that ProtBFN provides substantially better coverage of the protein space than the baseline methods. This result paired with Fig. 2f reinforces the idea that ProtBFN creates novel sequences that still cover the functional protein space present in UniProtKB.

To visualise the distribution coverage, we compute the embeddings of the protein sequences using a state-of-the-art protein language model and project these into two dimensions (Fig. 3). The close match between ProtBFN and UniProtCC’s visualised distributions highlights that the generated sequences provide broad coverage of the natural distribution even in this biologically meaningful representation space. Additional measures supporting ProtBFN’s extensive coverage of the protein space are detailed in Supplementary Section B.

**Figure 3.**
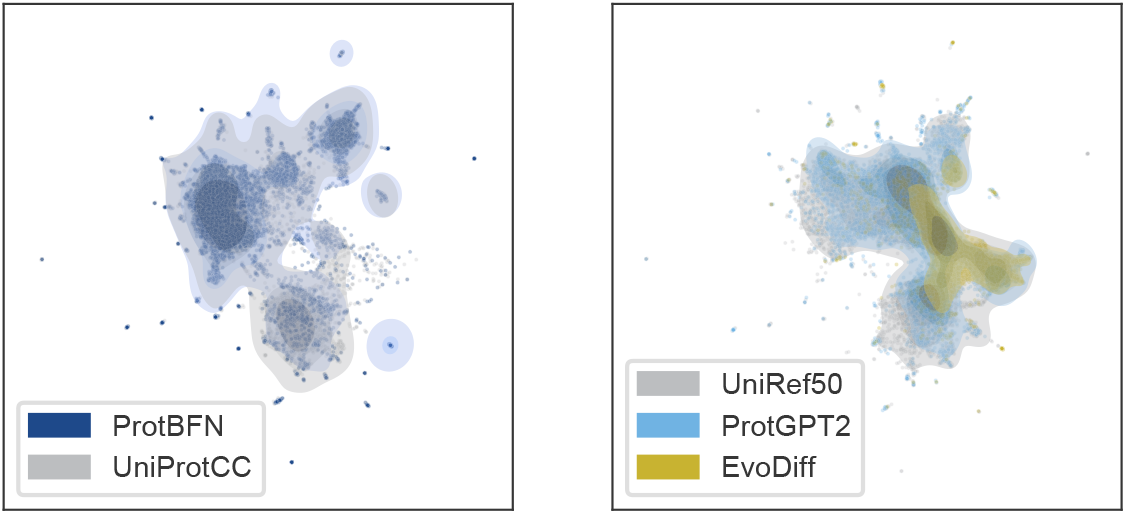
ProtBFN sequences show broad coverage of the training distribution in embedding space of a protein language model. To visualise distributions of protein sequences, the mean embedding of the ESM-2 model is calculated for 10 000 samples from each of ProtBFN, ProtGPT2 and EvoDiff and projected into two dimensions using the UMAP algorithm [39]. The projection is calculated using the union of both the UniProtCC and UniRef50 training distributions, and each method is overlaid with it’s respective training distribution.

### ProtBFN generates globular structural motifs with novel sequences

Protein function is closely tied to structure [4, 65]. Therefore, to characterise *de novo* protein sequences, which may differ substantially from those observed in nature, we analyze their correspondence with empirically determined protein conformations. The CATH S40 database [54] contains approximately 30 000 non-redundant, experimentally solved, protein domains that provide a broad coverage of known structural diversity. We compare the structural similarity of 2000 sequences sampled from each model against CATH S40 domains using template modeling (TM) scores. The TM1- and TM2-scores are normalised against the length of the generated protein and the length of the CATH S40 domain, respectively, with a TM score above 0.5 generally recognised as indicative of the same fold [69]. To avoid matching only fragments of CATH domains to generated proteins, or vice versa, a positive match requires both TM1 and TM2 scores to exceed 0.5.

ProtBFN acheives a CATH hit-rate of 65.7 %, surpassing ProtGPT2 (25.3 %) and EvoDiff (12.0 %). Moreover, these hits are of higher quality with 68.0 % of ProtBFN hits have sequential structure alignment program (SSAP) scores [44, 60] above 80 - which corresponds to the highest level of similarity (homologous superfamily) in CATH’s classification - compared to 38.0 % and 19.2 % for ProtGPT2 and EvoDiff, respectively (Fig. 4c). Furthermore, the proportion of near-complete hits, where both TM scores approach 1, (Fig. 4a), shows that ProtBFN generates longer sequences that fold into known domains more frequently. This is supported by the fact that ProtBFN samples retain high SSAP scores across all sampled protein lengths (see Supplementary Section C). However, despite high structural correspondence, ProtBFN’s sequences exhibit low sequence similarity to their CATH S40 targets (80.4 % of hits have under 50 % sequence similarity, Fig. 4b). This capability to produce recognisable globular folds with novel sequences is a prerequisite for the use of ProtBFN for rational protein design, and suggests a meaningful capability to go beyond training data and explore the uncharted proteome.

**Figure 4.**
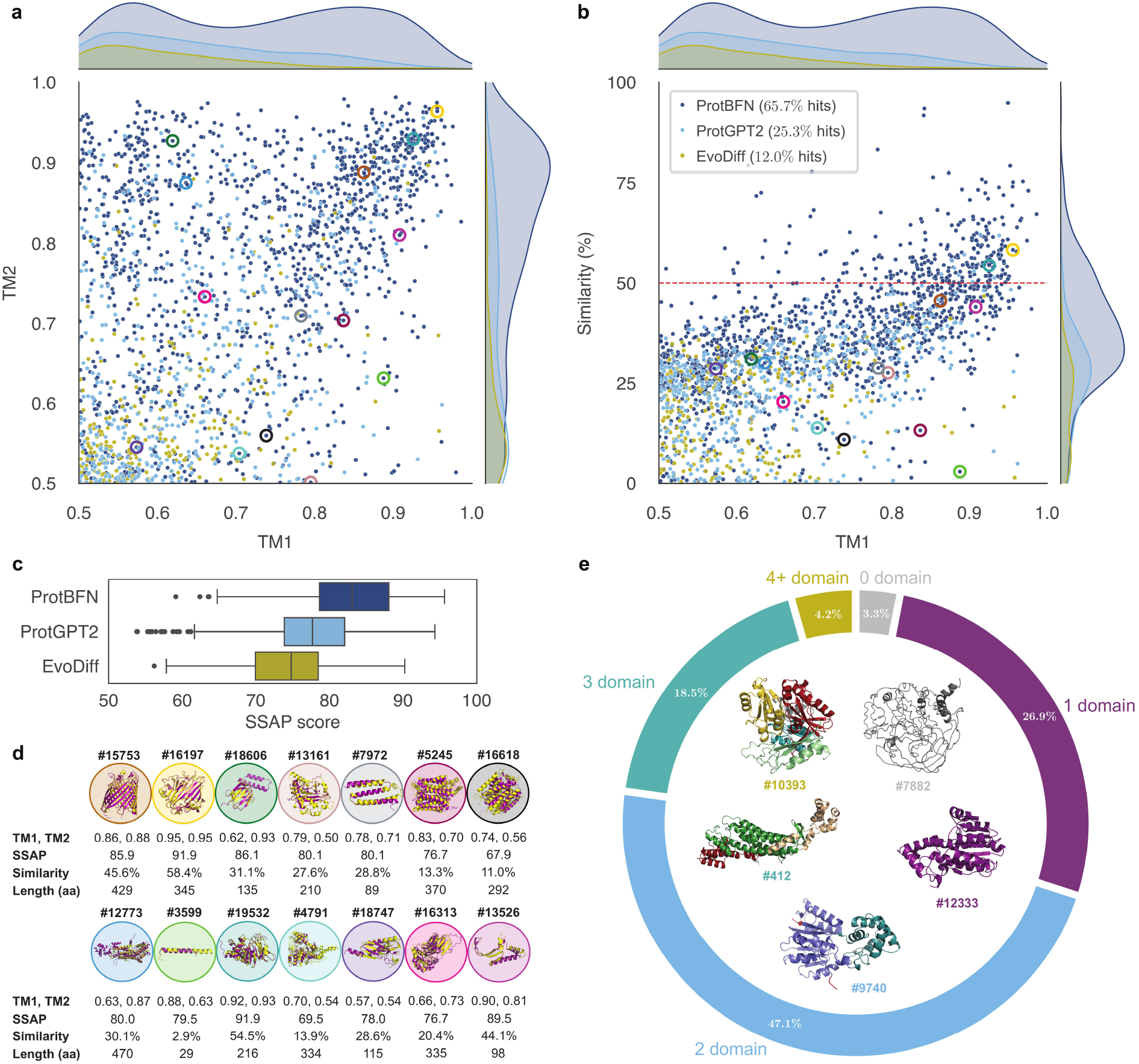
ProtBFN generates sequences that fold into naturally occuring structural motifs. Generated samples are folded with ESMFold, and the resulting structures are searched against the CATH S40 database, with a ‘hit’ determined by TM1 and TM2 scores above ProtBFN is found to generate hits significantly more often (65.7 % of samples) than the baseline ProtGPT2 (25.3 %) and EvoDiff (12.0 %) models. The distribution of TM1 and TM2 scores (**a**), and TM1 against sequence similarity (**b**) highlight the frequency with which ProtBFN also generates higher TM scores whilst maintaining novelty with respect to the reference sequence of a CATH domain. A more granular analysis of the structural hits its given by sequential structure alignment program (SSAP) scores [44, 60] (**c**). SSAP scores of 60–70, 70–80, and 80–100 correspond to similarity at the architecture, topology, and homologous superfamily levels, respectively, in the CATH nomenclature. ProtBFN hits scores higher than those of both baseline models, with vast majority showing similarity at the topological level or above. The highlighted elements on these scatter plots are detailed in **d** and selected to illustrate the diversity of structures and functional types generated by ProtBFN (a detailed discussion can be found in the main text). Each selected sample is annotated with sample number, TM1, SSAP, Similarity, Length and TM2. The sample structure is displayed in purple, while the CATH domain is displayed in yellow. Finally, Merizo-based domain segmentation of ProtBFN samples (**e**) reveals that zero-, single- and multi-domain samples are generated.

It is important to once again consider the diversity of the generated samples and, as illustrated in Fig. 4d, ProtBFN also excels in generating a structurally diverse array of functionally complex protein domains. The model spans the breadth of classes catalogued in CATH, including alpha helical (samples **7972, 5245, 16618, 12773, 3599**), beta sheet (**15753, 16197, 18606**), alpha-beta (**19532, 4791, 18747, 16313**), and irregular domains (**13525**). ProtBFN also effectively models various functional types, such as transmembrane proteins, including porins (**15753**) and transporters (**16618**), along with enzymes (**2773, 13161**). Additionally, the generated globular proteins span small (**18606, 7972, 18747**) and large (**16197, 5245, 4791**) structural domains, irrespective of CATH class.

Lastly, we used Merizo [34], a tool for domain segmentation, to understand the domain distributions of ProtBFN samples (Fig. 4e). Close to half of the tested samples have two domains, followed by single domain sequences with 26.9 %, and three or more domains with 22.7 %. Interestingly, the multi-domain constructs typically exhibit domain-domain interfaces, characterised by different types of interactions, rather than being simply connected by disordered loops (Supplementary Section C). This highlights that ProtBFN is able to generate proteins with globally coherent interactions, even when the interacting residues are separate in sequence space and belong to different folds. More broadly, this analysis demonstrates the structural and functional depth of ProtBFN’s learned distribution, highlighting the potential applicability of BFN generation across a diverse range of biological and biotechnological applications.

### AbBFN: A foundational sequence model tuned specifically for antibodies

The design of antibodies, a class of proteins that identifies and neutralises pathogens, is central to protein engineering applications such as monoclonal antibody-based cancer therapies [68]. To adapt our foundational generative model, ProtBFN, for specialised domains, we developed AbBFN, an antibody-specific model, by finetuning on variable heavy (VH) chains from the Observed Antibody Space (OAS) database [42]. We consider two validation sets in our analysis; 20 000 VH samples uniformly drawn from OAS and heavy-chain sequences from the SAbDab benchmark [33]. To ensure no data leakage, our training set is filtered of sequences similar to those in either validation set, leaving 195 M training examples. Full details on the training data and process are provided in Methods.

### Unconditional generation of antibody chains

Analogous to our investigation of ProtBFN, we validate the unconditional generation of AbBFN by comparing various metrics between the the generated and training (OAS) distribution. Detailed results in Supplementary Section D confirm that AbBFN accurately captures the natural distribution of VH chains.

### Zero-shot inpainting of antibody chains

Antibody VH chains are segmented into three complimentary determining regions (CDRs) – which primarily determine the binding specificity – and four framework (FR) regions – which provide a scaffold for the CDRs [5, 11] as well as providing important contributions to functions such as stability, including antibody-pairing and antibody expression [9, 37, 38, 57], as well as binding properties [45]. A common task for antibody sequence models is to conditionally generate subsets of these regions based on a given partial sequence (‘inpainting’), with performance measured in terms of the amino acide recovery (AAR) rate. Recalling that our BFN model can sample from the conditional distribution of any subset of variables (i.e. amino acids) despite being trained only for the unconditional generation - antibody inpainting represents a natural task with which to assess these zero-shot conditional generation capabilities.

We compare to leading antibody-specific language models AntiBERTy [51] and AbLang2 [43], which were trained for BERT-style conditional generation. As both of these methods were trained on the entire OAS, they have been exposed to both our validation datasets, potentially causing data leakage. Despite this, AbBFN is able to recover individual FR and CDR regions as well as these specialist models on the OAS validation set (Table 2), whilst demonstrating AAR rates consistent with the known increased variability of CDRs, and in particular CDR-H3. Noteably, AbBFN significantly outperforms the baseline models at predicting all FR regions simultaneously. We attribute this to larger masked region being out-of-distribution for BERT-style methods, which does not afflict the more flexible generation of a BFN.

**Table 2.**
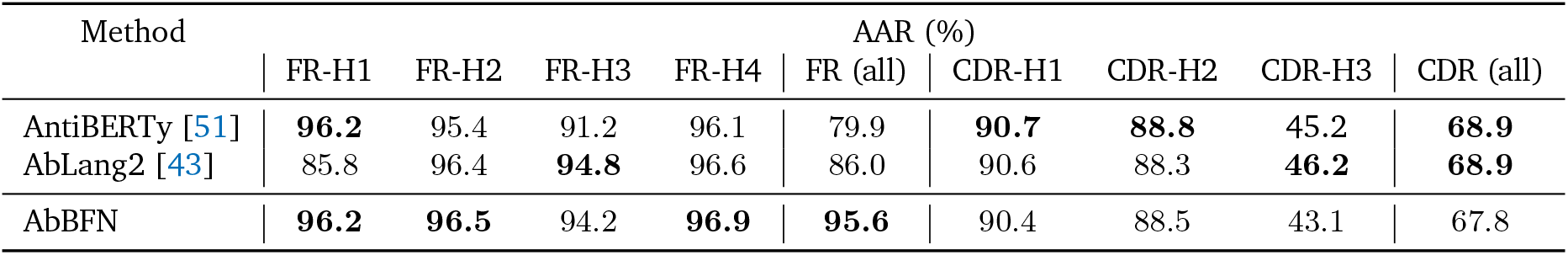
AbBFN can inpaint antibody heavy chain regions with zero-shot conditional generation. Amino Acid Recovery (AAR) rates on 20 000 VH chains sampled uniformly from OAS. Note that the training data of AbBFN was cleaned of similar sequences to this test set, whereas AntiBERTy and AbLang2 were trained on the complete OAS dataset and therefore have previously been exposed to these samples. Nevertheless, the zero-shot conditional generation of AbBFN has approximately the same, or improved, performance as these anti-body specific language models trained for inpainting tasks.

On the SAbDab benchmark (Table 3), AbBFN retains strong performance - significantly outperforming baseline methods which have not trained on this data, including those which can condition the generation on additional structural data [30, 33]. We attribute the increased gap between AbBFN and the language model methods to the removal of similar sequences to the SAbDab benchmark from our training data. To address this, we report the performance of AbBFN+, which underwent rapid fine-tuning (1000 adaptation steps) on the nine training folds for each test fold (see Methods). AbBFN+ outperforms all methods on CDR-H1 and CDR-H2 and significantly closes the gap on CDR-H3.

**Table 3.**
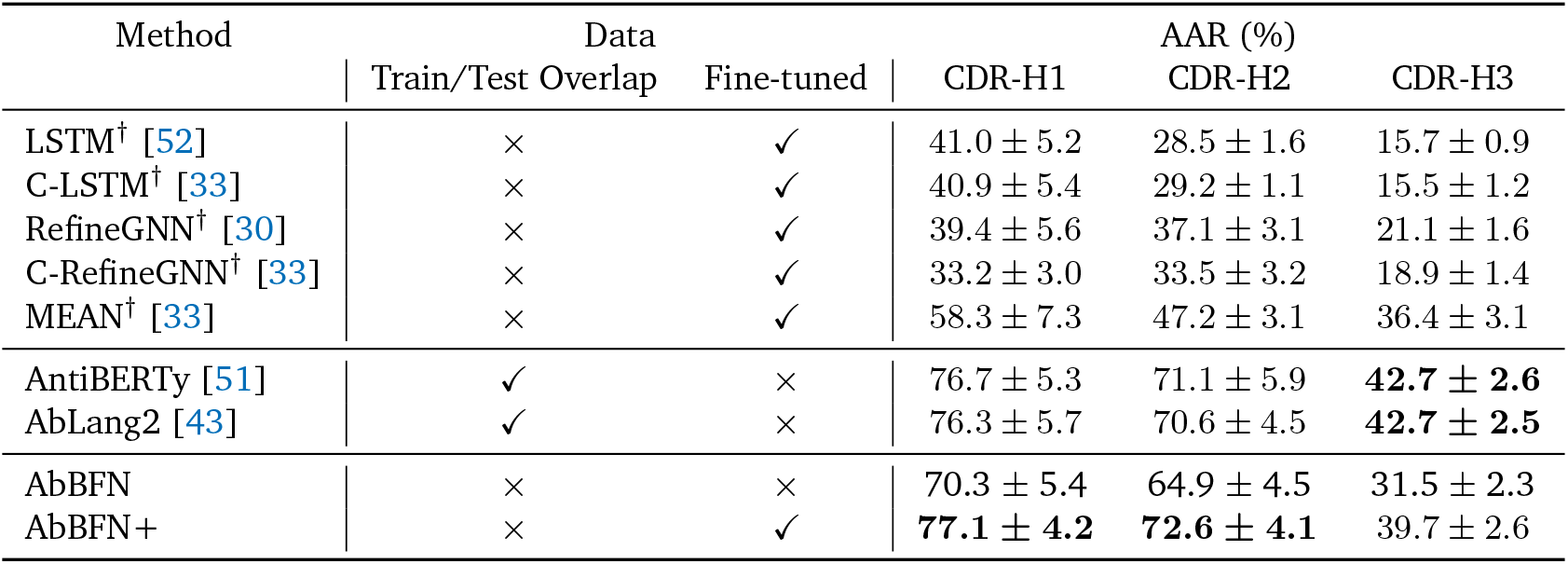
AbBFN baseline performance can be further improved with rapid fine-tuning. Amino Acid Recovery (AAR) rates for various methods on the 10-fold SAbDab benchmark. Methods labelled with the symbol † report results from [33]. ‘Train/Test Overlap’ report if a model’s pre-training data was cleaned to prevent data leakage into the SAbDab benchmark. AbBFN is found to provide leading performance among methods that avoid data leakage. For these models that have not seen the test distribution, ‘Fine-tuned’ reports if they were subsequently fine-tuned on the nine train folds associated with each test fold. AbBFN+ is obtained using 1000 steps of fine-tuning, and reports, or approaches, leading performance across all CDR regions.

| Method | Data |  | AAR (%) |  |  |
| --- | --- | --- | --- | --- | --- |
|  | Train/Test Overlap | Fine-tuned | CDR-H1 | CDR-H2 | CDR-H3 |
| LSTM<sup>†</sup> [52] | × | ✓ | 41.0 ± 5.2 | 28.5 ± 1.6 | 15.7 ± 0.9 |
| C-LSTM<sup>†</sup> [33] | × | ✓ | 40.9 ± 5.4 | 29.2 ± 1.1 | 15.5 ± 1.2 |
| RefineGNN<sup>†</sup> [30] | × | ✓ | 39.4 ± 5.6 | 37.1 ± 3.1 | 21.1 ± 1.6 |
| C-RefineGNN<sup>†</sup> [33] | × | ✓ | 33.2 ± 3.0 | 33.5 ± 3.2 | 18.9 ± 1.4 |
| MEAN<sup>†</sup> [33] | × | ✓ | 58.3 ± 7.3 | 47.2 ± 3.1 | 36.4 ± 3.1 |
| AntiBERTy [51] | ✓ | × | 76.7 ± 5.3 | 71.1 ± 5.9 | **42.7 ± 2.6** |
| AbLang2 [43] | ✓ | × | 76.3 ± 5.7 | 70.6 ± 4.5 | **42.7 ± 2.5** |
| AbBFN | × | × | 70.3 ± 5.4 | 64.9 ± 4.5 | 31.5 ± 2.3 |
| AbBFN+ | × | ✓ | **77.1 ± 4.2** | **72.6 ± 4.1** | 39.7 ± 2.6 |

## Discussion

In this work, we have demonstrated that BFNs have significant advantages in terms of both performance and flexibility compared to traditional methods for protein sequence modelling. ProtBFN generates plausible, diverse and novel protein sequences more effectively than leading autoregressive and discrete diffusion models. Additionally, AbBFN highlights that these models can be fine-tuned for domain specific capabilities, achieving or surpassing the conditional generation performance of BERT models that were trained specifically for sequence inpainting tasks. These capabilities underscore the potential of BFNs for the exploration of uncharted regions of the proteome and rational protein design.

Given the promise shown on protein generation, it appears reasonable to extend the application of BFNs to other biological sequence modelling tasks, such as RNA and DNA. More broadly, the ability of BFNs to handle different data modalities within a unified framework suggests the possibility of building multimodal models, for example that integrate sequences and structural data [47, 55]. Combined with the arbitrary zero-shot conditional generation we demonstrate in this paper, such multimodal BFN’s could provide unparalleled generative flexibility, further enhancing our ability to design and understand complex biological systems.

Nevertheless, it is important to acknowledge that BFNs are still an emerging technology with considerable fundamental questions remaining unexplored. In this direction, recent insights connecting BFNs with diffusion models thought stochastic differential equations [66], may facilitate the incorporation of the plethora of advanced sampling methods, e.g. [22, 26, 67], for diffusion, thereby providing even stronger performance.

## Acknowledgements

This research was supported with Cloud TPUs from Google’s TPU Research Cloud (TRC).

## Author contributions

TB and TA conceived the research, wrote the code, performed experiments and co-wrote the manuscript. LC assisted with design and implementation of biological validation, and in writing the manuscript. AG supported the work with Bayesian Flow Networks and developed modified sampling methods for conditional generation. BG and MG supported the validation of AbBFN. SC supported the validation of ProtBFN. LR curated and implemented custom training and validations datasets. AL supported interpreting results and assisted in the preparation of the manuscript. TB supervised the project.

## Data availability

All training data used in this work is available from publicly available servers – UniProt (www.uniprot.org) and the Observed Antibody Space (opig.stats.ox.ac.uk/webapps/oas).

## Code availability

Source code for this work is available at https://github.com/instadeepai/protein-sequence-bfn and model weights for ProtBFN and AbBFN can be accessed on Hugging Face at https://huggingface.co/InstaDeepAI/protein-sequence-bfn. Baseline results in Tables 2 and 3 use the available open-source implementations for AntiBERTy (https://github.com/jeffreyruffolo/AntiBERTy) and AbLang2 (https://github.com/oxpig/AbLang2).

## Computational resources

ProtBFN was pre-trained for approximately 2 weeks on a v4-256 TPU instance. AbBFN was fine-tuned from ProtBFN for approximately 4 days on a v4-128 TPU instance.

## Methods

### Bayesian Flow Networks for Discrete Data

#### Overview

Let *p*(**x**) be a distribution over *D* discrete variables that we wish to approximate. For some **x** ∼ *p*(·) we have that **x** = [*x*_1_, …, *x*_*D*_] ∈ *V* ^*D*^, where *V* = [*υ*_1_ … *υ*_*K*_] is a vocabulary with | *V*| = *K*. The BFN approximation to *p*(**x**) is constructed through the iterative revealing of information about **x** through a sequence of noisy observations **y**^(1:*i*)^ = {**y**^(1)^ … **y**^(*i*)^}.

A noisy observation at step 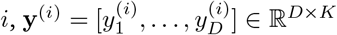 is drawn from the *sender distribution* as a normally-distributed vector for each variable,

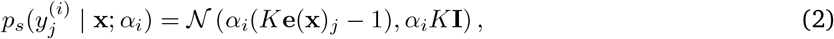

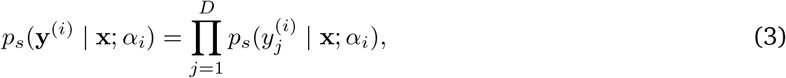

where *α*_*i*_ ∈ R^+^ is the *accuracy* at step *i*, such that a higher *α*_*i*_ reveals more information about the true value of **x**, and **e**(**x**) ∈ R^*D×K*^ is a one-hot encoding of **x**,

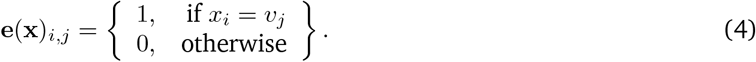

By starting from some prior distribution *θ*^(0)^ over the value of **x** and taking into account all observations so far, the *input distribution* is derived, providing the optimal estimation of the likelihood of each variable *x*_*j*_ independently,

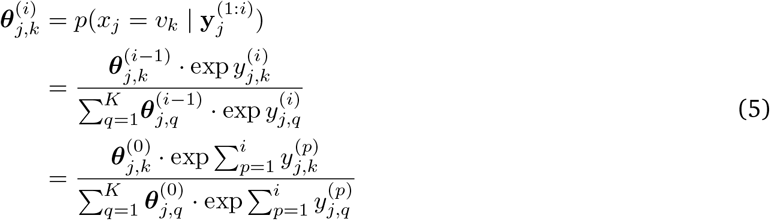

As *θ*^(*i*)^ is constructed based on an *independent* assumption about *x*_1_ … *x*_*D*_, it is clearly suboptimal whenever these variables are interdependent. The neural network *φ* is the *output distribution*, which aims to provide a better estimation by taking into account the interdepency between variables. Specifically,

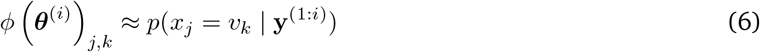

This approximation implies an equivalent approximation of the distribution of the *next* noisy observation **y**^(*i*+1)^. This approximation, referred to as the *receiver distribution*, is a mixture of Gaussians for each variable,

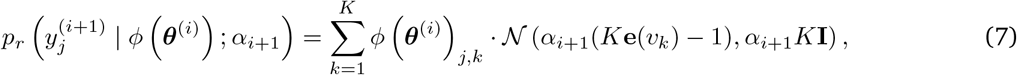

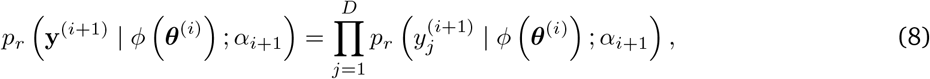

The error in *φ* is measured through the KL-divergence between *p*_*s*_ and *p*_*r*_ at each step. For *N* steps,

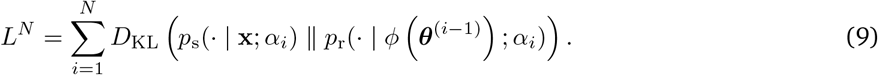

In practice, it is possible to derive a *continuous-time loss L*^*∞*^ where *N* → ∞. Under the assumption of a uniform prior *θ*^(0)^, the loss becomes remarkably simple and easy to approximate through Monte Carlo integration,

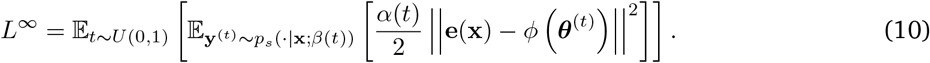

This loss function reflects the transition from an *N*-step process to a continuous time process, where *t* moves from 0 to 1. The sequence of accuracies *α*_1_ … *α*_*N*_ is replaced with the monotonically increasing *accuracy schedule*,

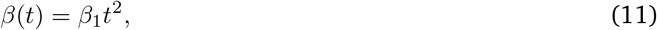

where *β*_1_ ∈ R^+^ is the final accuracy at *t* = 1, and its derivative scales the loss through time,

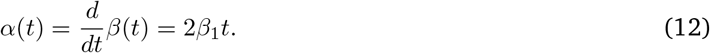

The continuous-time loss corresponds to a negative variational lower bound [23], and therefore by optimising the parameters of *φ* to minimise it, we arrive at an approximation of the true distribution *p*(**x**) from which **x** was drawn. To train our models, we follow the general procedure outlined in [23]. Specifically, the computation of the loss is described in Algorithm 1.

##### Algorithm 1

Continuous-time loss

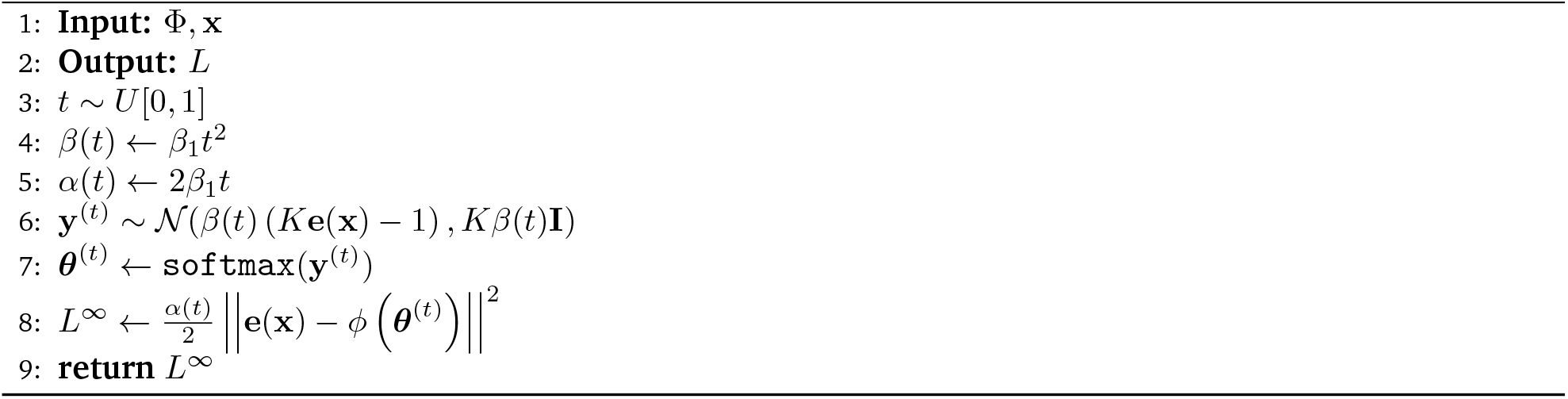

#### Entropy Encoding

In [23], the current time *t* is presented to the output network alongside the input distribution *θ*^(*t*)^. During initial sampling experiments, we discovered that, during the sampling process, the entropy of the input was noticeably higher at a given time *t* in comparison to that observed during training. We believe that this phenomenon occurs as the input distribution *θ*^(*t*)^ contains additional entropy from uncertainty in the output distribution (*φ θ*^(*t*)^). When time *t* is presented as an additional input to the network, this mismatch is, essentially, out of distribution for the output network, hampering performance. To resolve this, we replace the conventional Fourier encoding of time *t* used in [23] with a Fourier encoding of the entropy of each variable, appended to its corresponding input distribution before being passed into the network.

#### Sampling

We explored a variety of alternative sampling methods that reduce the overall temperature of sample generation from ProtBFN. In the conventional discrete-data sampling method described in [23], restricting ourselves to the logit space, the discrete sample generation process moves from **y**^(*t*)^ at time *t* to **y**^(*s*)^ at time *s* according to the equation,

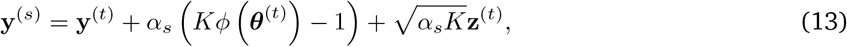

where *θ*^(*t*)^ = `softmax`(**y**_*t*_), **z**^(*t*)^ *∼ 𝒩* (**0, I**) is isotropic noise and *α*_*s*_ = *β*(*s*) *− β*(*t*) is the change in accuracy. By the additivity of normally distributed variables and with **y**^(0)^ = **0**, the distribution of **y**^(*s*)^ given preceding steps at times *i* = 0, … *t* is,

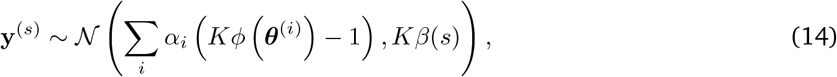

or equivalently,

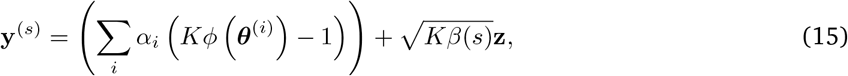

where **z** ∼ *N*(**0, I**) is a fixed isotropic noise. Sampling according to Equation 15 would yield an ODE similar to that described in [66]. However, we observed that simply by replacing the summation of previous predictions with the most recent prediction, e.g.,

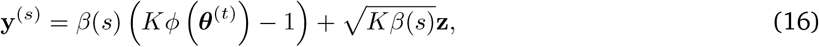

##### Algorithm 2

Reconstructed ODE Sampling

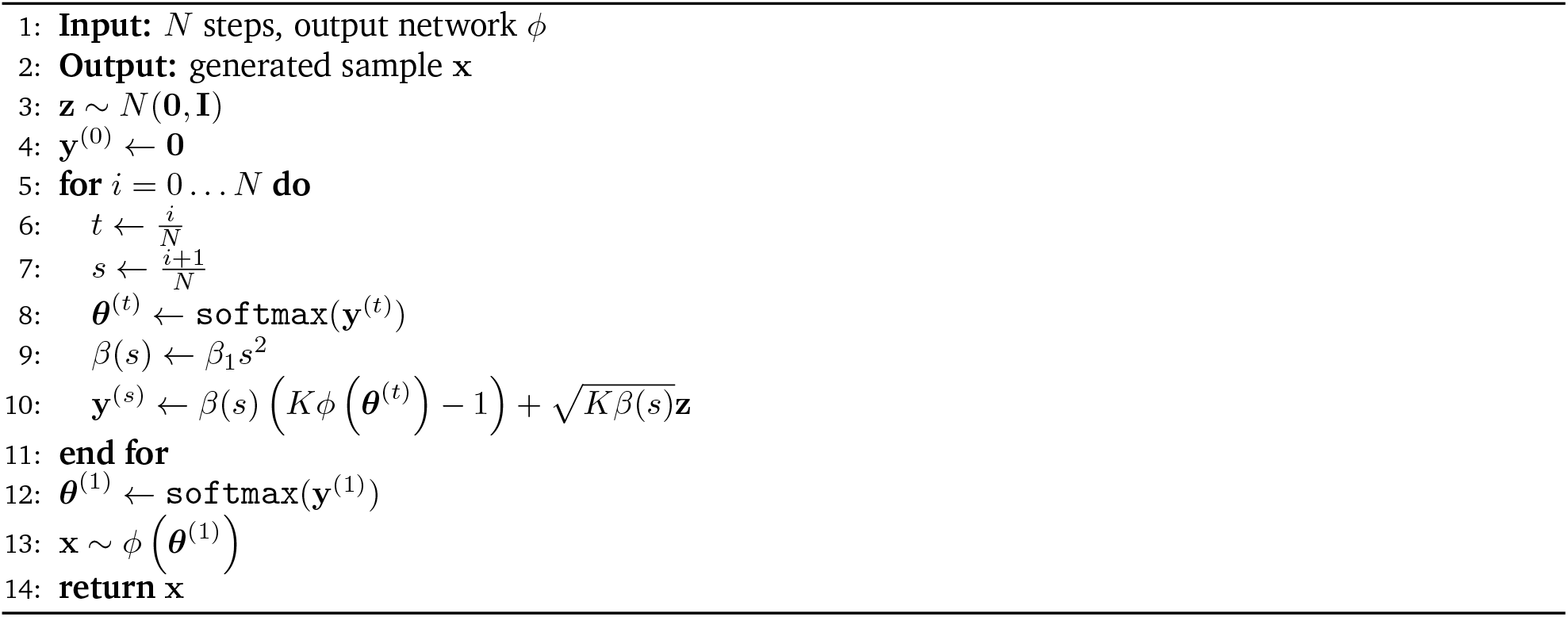

substantially reduced the perplexity of generated samples. Effectively, this method is equivalent to taking our most recent prediction *φ* (*θ*^(*t*)^) and supposing that we had predicted it at every step 0 … *s*. This is a similar method to that used in [47], except that the direction of the noise remains stationary throughout the sampling process, as we found this ‘ODE-like’ behaviour to be a more stable approach. Our overall sampling method is described in Algorithm 2.

### Inpainting

Consider the task of inpainting some sequence **x** according to a binary mask **m** ∈ [0, 1]^*D*^ where **m**_*i*_ = 0 indicates that the *i*th element of **x** should be used as conditioning information, and **m**_*i*_ = 1 indicates that the *i*th element of **x** should be inpainted. Inspired by SMCDiff [62], we treat conditional generation of the inpainted regions as a sequential Monte Carlo problem which we may solve with particle filtering [15, 16]. Given an output network *φ* and input distribution *θ*^(*t*)^ at time *t*, the KL divergence of the sender and receiver distributions, with accuracy *α*, for the conditioning subset **x**_**m**_ is,

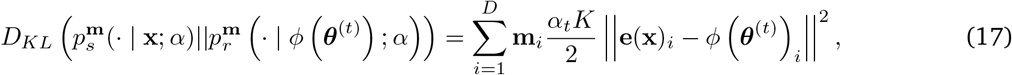

which we will denote *D*(**x, m**, *θ*^(*t*)^, *α*). Given *q* particles at time *t* with input distributions 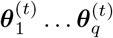, we resample each particle with probability proportional to 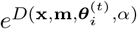. Combining SMC with our sampling method, we arrive at Algorithm 3.

### Pre-training ProtBFN

#### Model

ProtBFN is based on the 650-million parameter architecture used in [35] which is a BERT-style encoder-only transformer [12] with rotary positional embeddings [58]. The architecture consists of 33 layers, each consisting of 20-head multi-head self-attention [63] in 1280-dimensional space followed by a single-layer MLP with GeLU activation [24] and a hidden dimension of 5120. The only substantial difference between ProtBFN’s architecture architecture and that used in [35] is that the initial token embedding is replaced with a linear projection, as our network’s input *θ*^(*t*)^ is a distribution over possible token values.

##### Algorithm 3

Greedy SMC inpainting

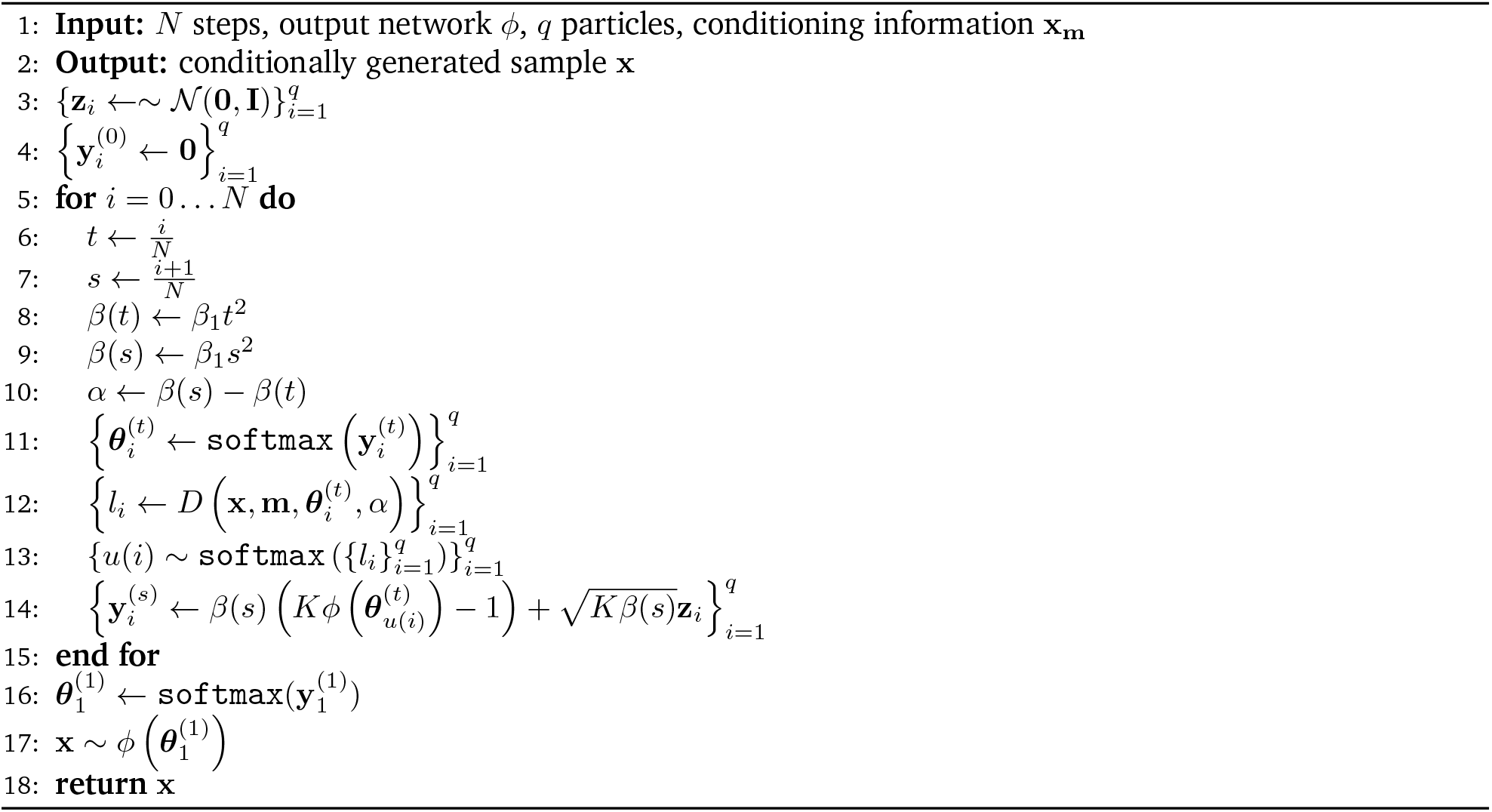

#### Data

ProtBFN is trained on data obtained from the January 2024 release of UniProtKB [1]. The data is filtered according to the ‘Protein Existence’ (*PE*) property, including only those proteins which are inferred from homology (*PE* = 3), has evidence at the transcript level (*PE* = 2) or evidence at the protein level (*PE* = 1). By removing those proteins which are known to be hypothetical (*PE* = 4) or are of unknown existence (*PE* = 5), ProtBFN is restricted to model the distribution of proteins that are very likely to exist, meaning that greater confidence can be placed in the sequences it generates. Additionally, we found that it was substantially faster to train a model when removing hypothetical proteins, indicating that they introduce substantial amounts of additional entropy which may imply their generally lower quality. Additionally, ProtBFN is trained only on those sequences with length *l* < 512, with the final token used to encode an end-of-sequence (EOS) token. After filtering by PE and length, the final training set contains 71 million sequences. Where clusters are used to reweight and debias the data, the clusters are obtained from UniRef50 [59]. Each sequence is represented by its amino acids followed by an end-of-sequence (EOS) token. All other tokens after EOS are PAD tokens which are treated as normal tokens for the purposes of noisy observations, predictions and the loss. As this dataset is a (C)leaned and (C)lustered subset of UniProt, we will refer to it for the rest of the text as UniProtCC.

#### Training

ProtBFN is first pre-trained for 250 000 training steps with a batch size of 8,192. We observed that a large batch size was necessary to obtain stable gradient estimates. Adam [32] is used with *β*_1_ = 0.9 and *β*_2_ = 0.98 as in [23]. The learning rate is initialised to 0 and linearly increased to 10^*−*4^ at step 10, 000, after which it is held constant. Throughout training, the norm of the gradient is clipped to 500. A copy of the network’s parameters is maintained with an exponential moving average of the weights with decay rate 0.999. During this first phase of training, samples are drawn uniformly at random from all training data.

Next, the model is trained for a further 250 000 steps with clustered data. Specifically, each cluster is constructed by taking all samples within the corresponding Uniref50 cluster which pass through both the PE and the length filters. During this training phase, each cluster is sampled with probability proportional to the square root of its size, so that ProtBFN is debiased away from those proteins most heavily studied by humans, but not overly focused on very data-sparse 1-member clusters as would happen when uniformly sampling clusters or training only on the Uniref50 cluster centers as done in [19]. Once a cluster has been sampled, any sequence contained within it is chosen uniformly at random. During this second phase of training, the optimiser is completely reset and again, the learning rate is linearly increased from 0 to 10^*−*5^ over the first 10, 000 steps. The main focus of this article is providing evidence BFNs can be used in the protein space. For more in-depth analysis concerning biases present in protein data please refer to [53], [43] and [14].

The training curve of ProtBFN is shown in Fig. 5. The loss decreases monotonically over the first 250,000 steps. At the point at which the cluster sample weighting is introduced, the loss increases significantly, reflecting the distribution shift of the underlying training data. The loss continues to monotonically decrease over the next 250,000 steps, although overall loss remains higher than during the initial training stage, indicating that the introduction of the weighted cluster sampling leads to a substantially higher underlying entropy of training data.

**Figure 5.**
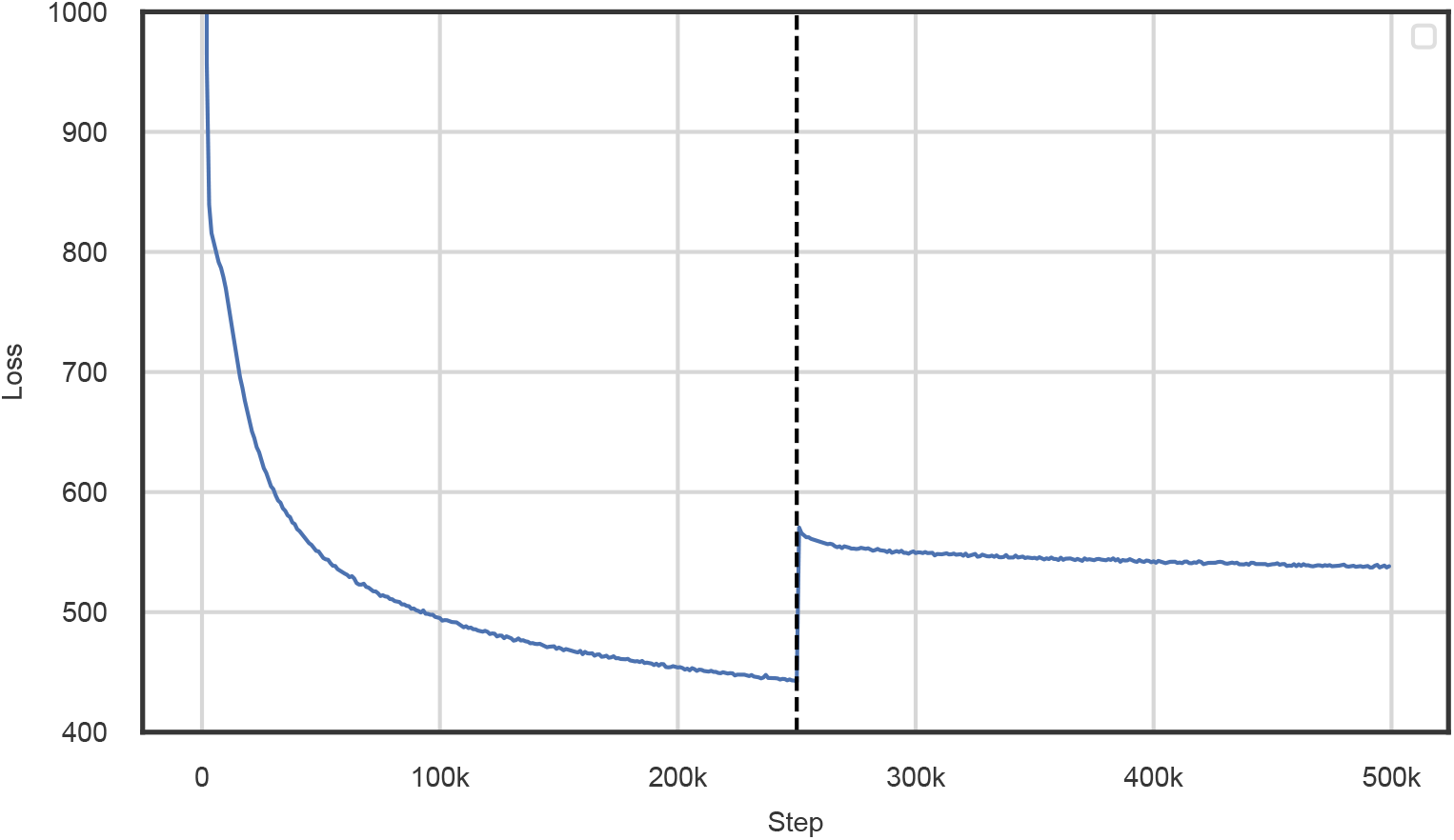
Training curve for the pre-training of ProtBFN. The loss is averaged over every 1000 consecutive steps to produce a smooth plot. The dashed line indicates the step at which the weighting of clusters is changed.

#### Sampling and Filtering

To generate the 10 000 samples used for ProtBFN *de novo* generation results, we use the sampling algorithm described in Algorithm 2 with *N* = 10000 e.g. 10000 sampling steps, and with the weights of the model obtained from the exponential moving average. We found that the lower-temperature sampling method occasionally produced pathological sequences that are highly repetitive. To remedy this, we counted the number of sequential repeats of any given amino acid and consider any amino acid ‘repetitive’ if it repeats more than 3 times. For any given sequence, the ‘repetitivity score’ was calculated as the total number of repetitive amino acids, divided by the sequence length. We discarded any sequence within the top 20th percentile with respect to their repetitivity score. Additionally, we found that the sampling method occasionally generated sequences with very high perplexity. We therefore discarded any sequence within the top 30th percentile with respect to their perplexity. This process rejected approximately 44% of generated samples.

#### Coverage Score

For a given number of sequences, we can estimate the expected number of unique clusters which the sequences would be assigned to. This estimate assumes that a given sequence can be assigned to multiple clusters, which is the case when clustering according to 50% sequence identity. If we sample *n* sequences randomly and independently from a set of clusters with normalised weighting *ω*_*i*_, the expected number of unique clusters sampled can be calculated as

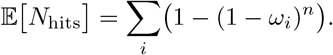

For ProtGPT2 and EvoDiff we use an equal weight for each cluster, since that corresponds to the UniRef50 dataset, while for ProtBFN we use cluster weights inversely proportional to the cluster size, as that corresponds to our training data. For our diversity score, we divide the number of unique clusters hit by the expected number of cluster hits as calculated above. This estimate does over-count expected number of unique clusters, since it treats the clusters independently; however, we compensate for that by normalizing the diversity scores by the equivalent metric calculated on a large subsample of our training data.

### Fine-tuning AbBFN

#### Data

We downloaded the heavy chain unpaired OAS dataset [42] on 21^*st*^ February 2024. We filtered the data with a similar procedure to [6], which is as follows:

1. Filter out the studies: “Bonsignori et al., 2016”, “Halliley et al., 2015”, “Thornqvist et al., 2018”.
2. Filter out studies originating from immature B cells (BType is “Immature-B-Cells” or “Pre-B-Cells”).
3. Filter out studies originating from B cell cancers (Disease is “Light Chain Amyloidosis” or “CLL”).

Then for each remaining study filter the sequences as follows:

1. Filter if: sequence contains a stop codon; sequence is non-productive; V and J regions are out of frame; Framework region 2 is missing; Framework region 3 is missing; CDR3 is longer than 37 amino acids; J region identity is less than 50%; CDR edge is missing an amino acid; locus does not match chain type.
2. Remove sequences if they only appear once in a study, then make unique.
3. Filter if the conserved cysteine residue is not present, or misnumbered in the ANARCI numbering.
4. Create the near-full-length sequence IMGT positions 21 through 128 (127 for light chains and for heavy chains from rabbits and camels). Filter if framework 1 region is < 21 amino acids.
5. Remove duplicate near-full-length sequences and tally up the counts, filter out sequences which only appeared once dropped on the grounds of insufficient evidence of genuine biological sequence as opposed to sequencing errors, following [6].
6. Filter out sequences which contain any amino acids which are not the standard 20.
7. We use the sequence from the full ANARCI [18] numbering, not the near-full-length sequence as in [6].

This corresponds to all of the filters/preprocessing described in [6] but using full variable domain sequence (insofar as it was present in the original sequence) instead of the near-full-length sequence. We applied one additional filter which removed any sequences which had an ANARCI numbering with an empty region.

To create the SAbDab test set we used SAbDab data downloaded on 29^*th*^ February 2024, specifically the summary table to select only the paired chains, ensuring we exclude single-domain antibodies; most of these are of camelid origin and therefore belong to germline genes which were not present in our training set. We also removed single-chain fragment variable (scFv) antibodies. We then parse SEQRES attributes from the original SAbDab PDB files, and ran ANARCI [18] on these using the IMGT numbering scheme to obtain the final variable domain sequences as ANARCI retains only the subset of the sequence that could be numbered, thereby removing additions such as purification tags.

To ensure that the training data is dis-similar to the testing data we split the separated OAS and SAbDab data into heavy and light chains. We then uniformly selected and set aside 20,000 heavy chain sequences from the OAS data at random and added all heavy chains from SAbDab to construct the test data set. Next, we used MMSeqs2 [56] to query this combined data set against the remainder of the OAS data set with default sensitivity of 5.7, minimum sequence identity of 0.95, coverage of 0.8 and coverage mode 0. We removed any hits from the training data. Due to the presence of heavily engineered antibodies, the test data originating from SAbDab is more out-of-distribution when compared to the test data originating from OAS and filtering out the similar sequences still retained 99% of the data, whereas filtering out sequences which are similar to the 20,000 uniformly-sampled test set sequences from OAS retained 79% of the OAS data. Filtering out these similar sequences produced 195M training examples, compared to 248M before sequence similarity filtering.

#### Training

AbBFN is fine-tuned from the ProtBFN model on the filtered OAS data. For computational efficiency, the maximum sequence length is reduced to 256. It is trained for 100,000 steps with a batch size of 8192. Adam [32] is used with *β*_1_ = 0.9 and *β*_2_ = 0.98 as in [23]. The learning rate is initialised to 0 and linearly increased to 10^*−*5^ at step 10, 000, after which it is held constant. Throughout training, the norm of the gradient is clipped to 500. A copy of the network’s parameters is maintained with an exponential moving average of the weights with decay rate 0.999. The training curve of AbBFN is shown in Fig. 6.

**Figure 6.**
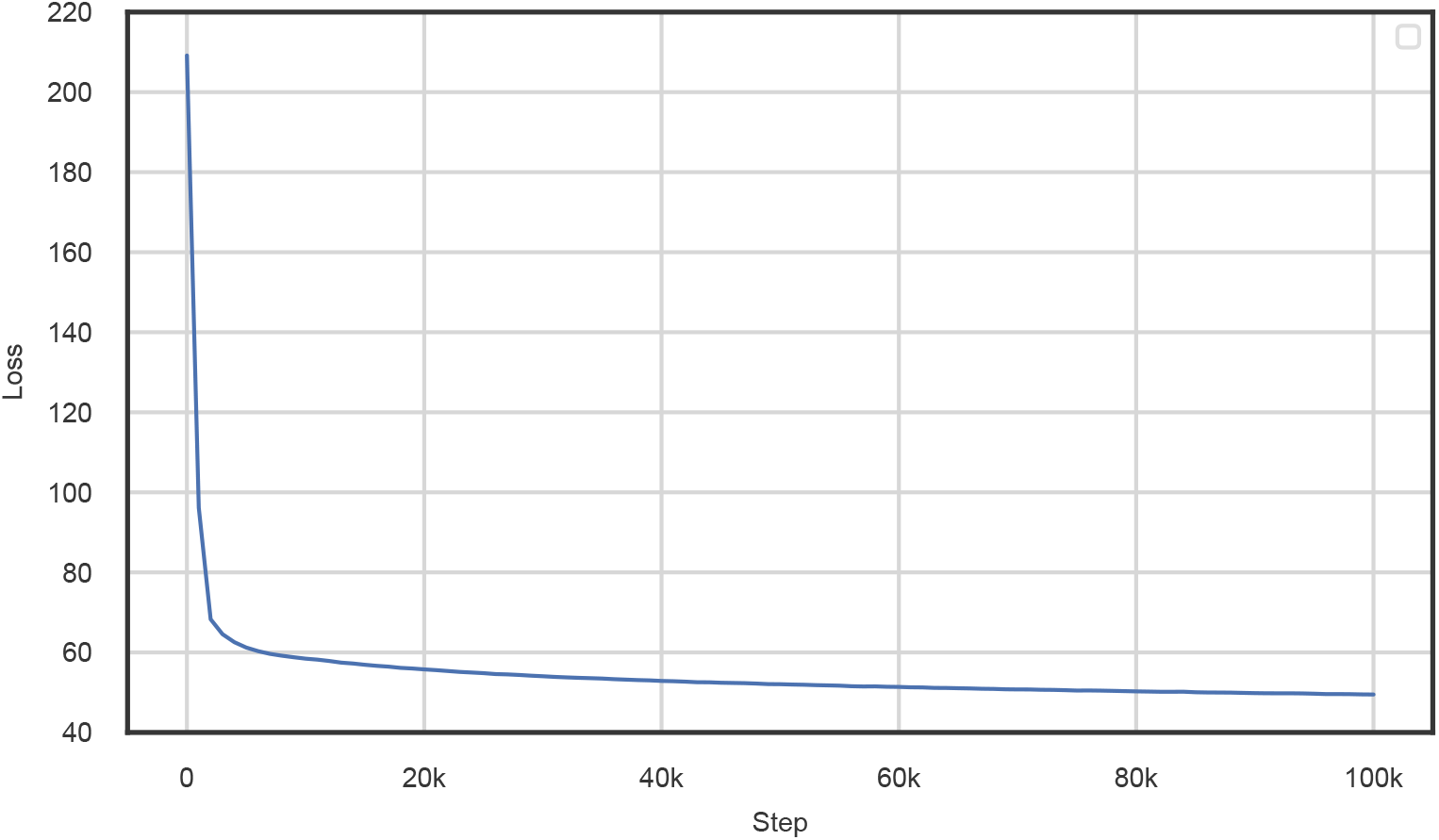
Training curve for the fine-tuning of AbBFN. The loss is averaged over every 1000 consecutive steps to produce a smooth plot.

#### Fine-tuning AbBFN+

To assess if AbBFN is able to learn the distribution of heavy chain sequences in SAbDab, we further fine-tune the model on each of the 9 train folds from the 10-fold cross validation splits used by [33], using the remaining test fold for evaluation. The model is fine-tuned for 1000 training-steps with a batch size of 512. Adam [32] is used with *β*_1_ = 0.9 and *β*_2_ = 0.98 and the learning rate is linearly increased from 0 to 10^*−*5^ over the 1000 training steps. Throughout training, the norm of the gradient is clipped to 500. A copy of the network’s parameters is maintained with an exponential moving average of the weights with decay rate 0.995. The exponential moving average parameters are used for zero-shot inpainting results.

#### Sampling AbBFN

To generate the 10 000 samples used for AbBFN *de novo* generation results, we use the sampling algorithm described in Algorithm 2 with *N* = 10000 e.g. 10000 sampling steps, and with the weights of the model obtained from the exponential moving average. We did not find that it was necessary to filter samples by repetitiveness or perplexity, possibly due to the increased simplicity of the OAS domain in comparison to UniProtCC by virtue of the common immunoglobulin fold.

#### Inpainting

To inpaint OAS and SAbDab test sequences, we use the algorithm described in Section 3. We use *p* = 1024 particles for each sequence, and *N* = 100 sampling steps. For both AbBFN and AbBFN+, we use the weights of the model obtained from the exponential moving average during training.

