## Supplementary Information for "Protein Sequence Modelling with Bayesian Flow Networks"

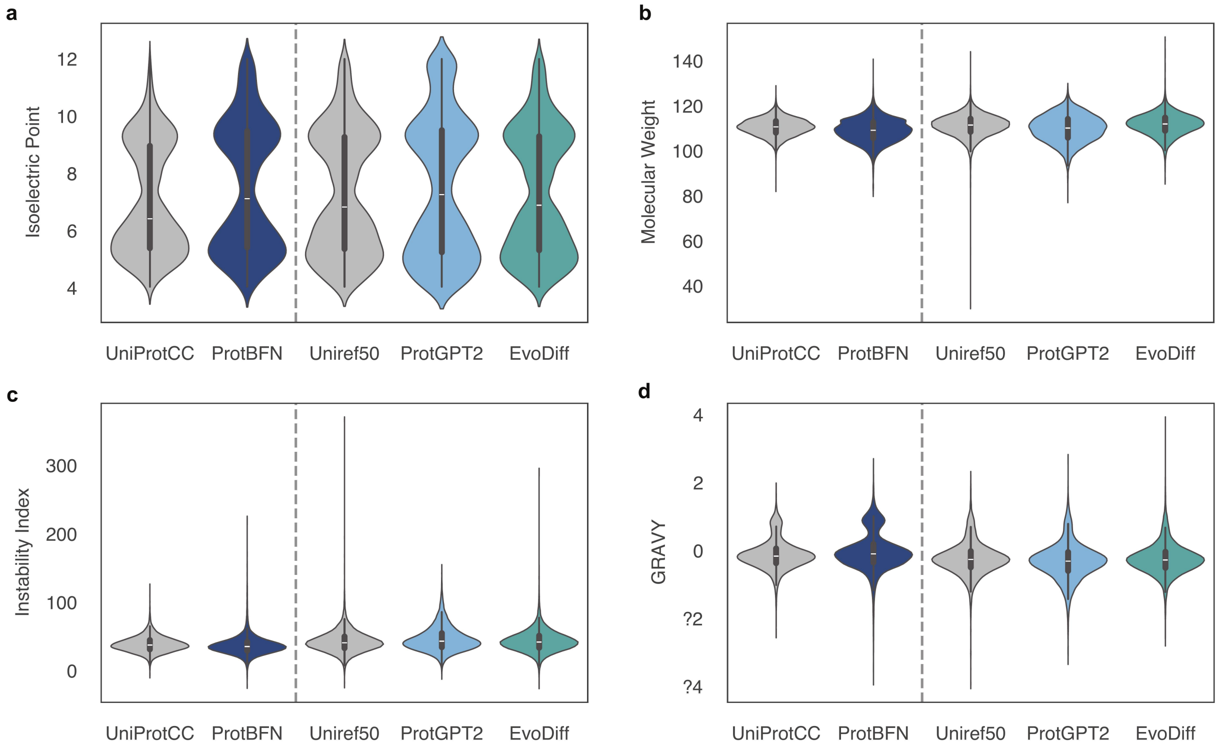


**Figure 1 | ProtBFN effectively *de novo* generates protein sequences that are in distribution with respect to various biochemical properties.** (**a**) the isoelectric point is the pH at which a given protein has a charge of zero, and typically represents represents the pH at which solubility is minimal. (**b**) the molecular weight of a given protein sequence can be used for further assessment of many functionalities, such as gene regulation and immunological responses. (**c**) the instability index of a protein sequence is an estimate for the likelihood that it will be stable in a test tube. (**d**) the **gr**and **av**erage of hydropathicit**y** (GRAVY) is the average hydropathy value of a protein, where negative GRAVY values are hydrophilic and positive values mean they are hydrophobic. In all cases, ProtBFN is seen to well match the natural training distribution UniProtCC, whereas ProtGPT2 and EvoDiff are substantially closer to UniRef50.

### Supplementary Information

#### A. Additional naturalness measures for generated proteins

**ProtParams** We compute a number of additional biochemical properties of naturalness of *de novo* generated proteins using BioPython [2], as shown in Fig. 1. Consistent with our other findings, we observe that ProtBFN is highly consistent with UniProtCC, whereas ProtGPT2 and EvoDiff are instead consistent with Uniref50. As UniProtCC is restricted to consider only those proteins observed in nature, the high coherence with ProtBFN implies that our generated *de novo* proteins are highly natural with respect to these important properties.

**NetSurfP-3.0** We compute a number of additional structural measures of naturalness of *de novo* generated proteins using NetSurfP-3.0 [6]. These are shown in Fig. 2. Each residue in each protein is annotated as one of 8 possible secondary structures, or intrinsically disoredered. Once again, we observe that ProtBFN is highly consistent with UniProtCC, whereas ProtGPT2 is instead consistent with Uniref50. We note that EvoDiff substantially under-represents a number of secondary structure types, particularly those which occur with lower frequency within Uniref50.

#### B. Additional coverage measures for generated proteins

**ESM-MMD estimator** Maximum mean discrepancy (MMD) is a metric defined over a space of probability distributions. It measures how different any two distributions are by embedding their support into a reproducing

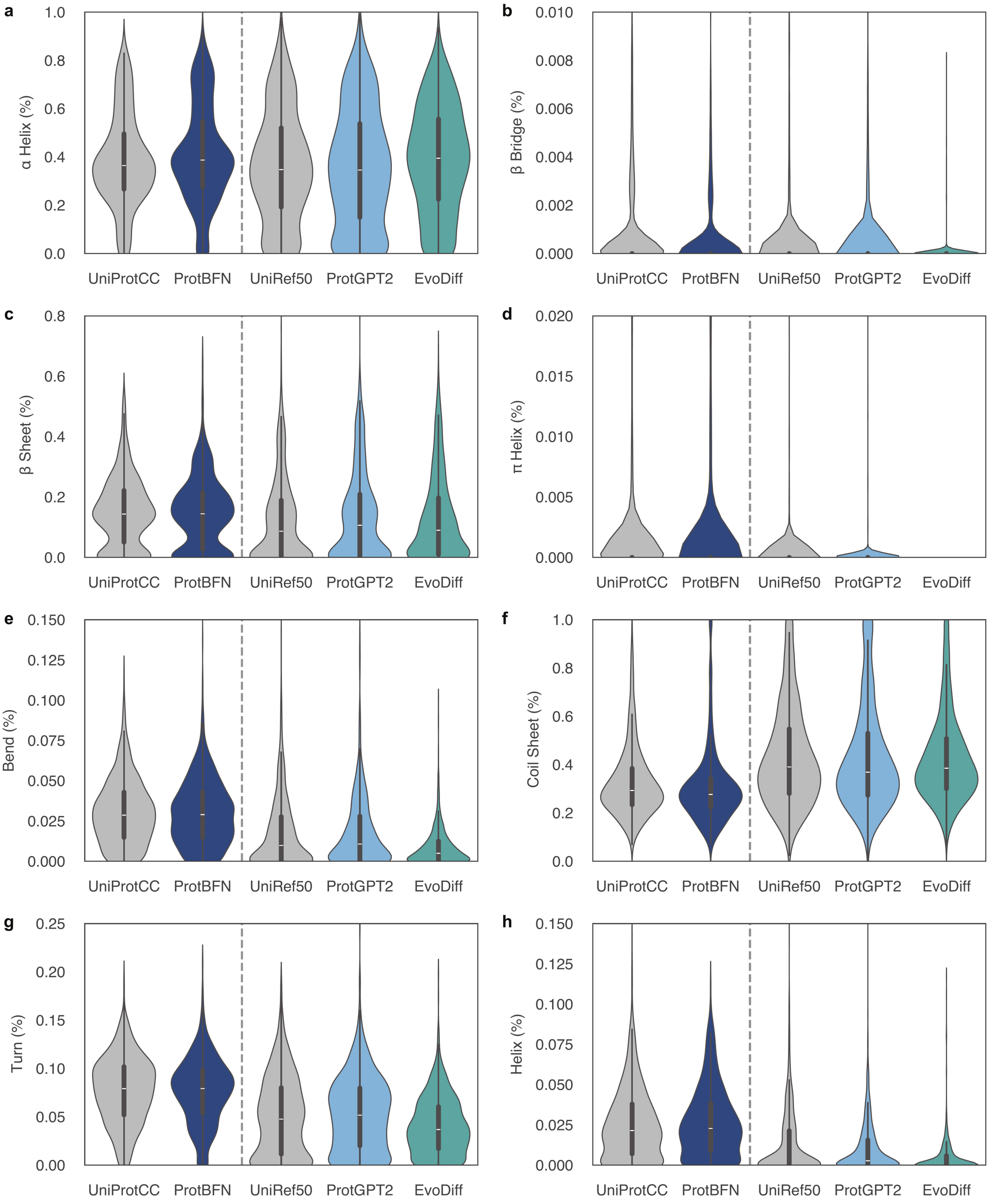


**Figure 2 | ProtBFN effectively *de novo* generates protein sequences that are in distribution with respect to various per-residue structural annotations.** Each plot shows the distribution of different secondary structures, as predicted by NetSurfP-3.0 [6], measured as a fraction of each overall protein's residues. In all cases, ProtBFN is seen to well match the natural training distribution UniProtCC, whereas ProtGPT2 and EvoDiff are substantially closer to UniRef50.

kernel Hilbert space. An unbiased estimator for the squared MMD can be calculated using a kernel as follows [5]

$$\widehat{\mathrm{MMD}}^2 = \frac{1}{N(N-1)} \sum_{i \neq j} \big( k(x_i, x_j) + k(y_i, y_j) - 2k(x_i, y_j) \big),$$

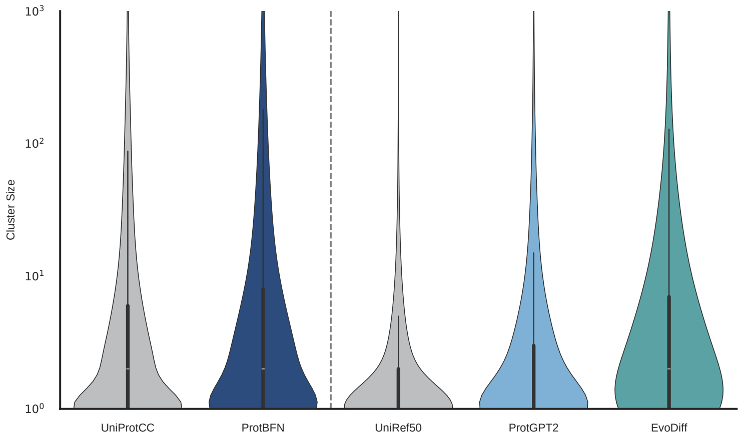


**Figure 3** | Sizes of unique clusters hit

where $\{x_i\}_{i=1}^N$ and $\{y_i\}_{i=1}^N$ are i.i.d. samples from each distribution, and $k$ is a symmetric positive-definite kernel function. We use the radial basis function kernel to estimate the squared MMD between mean ESM embeddings of each sequence dataset. Table 1 shows the estimated $\mathrm{MMD}^2$ between each set of sequences. Note that this estimator is unbiased and so can be negative, despite estimating a squared quantity. We observe that ProtBFN is substantially closer to UniProtCC than any other dataset, even including UniRef50. In contrast, ProtGPT2 is the closest model to UniRef50.

| Source | Target | $\widehat{\mathrm{MMD}}^2$ ($\times 10^{-3}$) | | |
|---|---|---|---|---|
| | | Mean | Median | Spread |
| UniProtCC | UniProtCC | -0.295 | -0.295 | $\pm$ 0.034 |
| | **ProtBFN** | **6.512** | **6.512** | $\pm$ 0.031 |
| | ProtGPT2 | 18.832 | 18.832 | $\pm$ 0.017 |
| | EvoDiff | 62.941 | 62.940 | $\pm$ 0.014 |
| UniRef50 | UniRef50 | -0.277 | -0.277 | $\pm$ 0.028 |
| | UniProtCC | 10.555 | 10.555 | $\pm$ 0.019 |
| | ProtBFN | 16.487 | 16.487 | $\pm$ 0.015 |
| | **ProtGPT2** | **4.076** | **4.076** | $\pm$ 0.025 |
| | EvoDiff | 32.848 | 32.847 | $\pm$ 0.027 |

**Table 1** | $\mathrm{MMD}^2$ estimates between embeddings

**Cluster size coverage** We compare the distribution of the sizes of UniRef50 clusters covered by each model. We run `mmseqs2` against the UniRef50 cluster centers and record all clusters to which at least one generated sequence matches. We then take all of these clusters and plot the distribution of the sizes of these clusters as defined by UniRef50. UniRef50 has many clusters of size 1, and many of the sequences we removed when constructing our training data fall into these isolated clusters. For each value of cluster size, this metric provides an estimate of the number of UniRef50 clusters of that size which lie within the support of the model's learned distribution. Figure 3 shows the distribution of cluster sizes for each set of sequences. Since we consider only unique clusters, the distributions shown are not the relative frequency of the model sampling clusters of a given size, but rather the relative coverage of clusters of a given size. In other words, if a given model tends to sample a specific cluster more frequently than others, that is not shown here; what is shown, is the portion of clusters of each size that the model was able to sample from. A closer match between a model and its training dataset would indicate that sequences generated by the model have a broad coverage of the modes present in the dataset. Both ProtBFN and ProtGPT2 show good match in cluster size distribution to their respective training datasets, whilst the sequences from EvoDiff which did match clusters seem to over-represent larger clusters.

#### C. Additional structural analysis of generated proteins

**The sequential structure alignment program** SSAP is a metric used to generate the CATH hierarchical classification (Class, Architecture, Topology, Homology) and it calculates the pairwise similarity of structures. When comparing two structures SSAP takes into account beta carbon vectors, the order of residues and secondary structure elements. The algorithm returns a normalised score between 0-100. Pairs of proteins with a score bellow 60 typically don't show any degree of similarity. Scores between 60-70 correspond to the Architecture level in CATH, and usually have similar local structural elements and belong to the same protein class. Scores between 70-80 correspond to the Topology level in CATH and structures with these scores usually present the same overall fold but with variations in coils and other secondary structure elements. Lastly, scores between 80-100 correspond to the Homologous Superfamily level in CATH, and structures with these scores display extremely similar overalls folds [8].

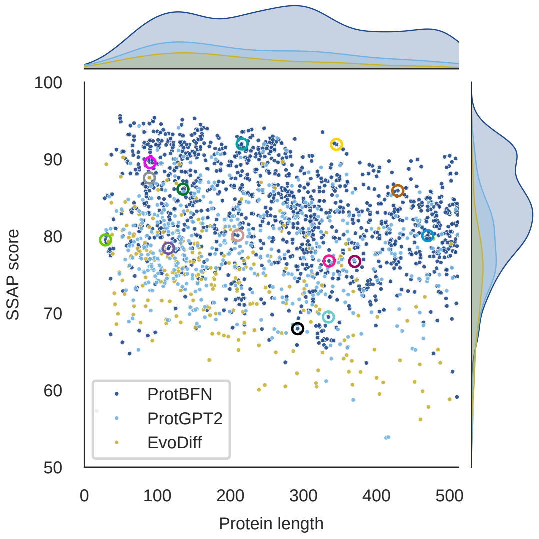


**Figure 4** | **Sample length vs SSAP score for ProtBFN (dark blue), ProtGPT2 (light blue) and EvoDiff (yellow).** The SSAP score is calculated between the sample and the best CATH hit.

**Interacting domains** We used the Merizo algorithm to segment BFN samples into individual domains. For our multi-domain samples we wanted to understand if they are simply chained together by flexible loops or if there is some type of global coherence. To this end we used a popular method, called getcontacts [4], to calculate the number of interactions between domains annoted by Merizo. The interactions considered are: hydrogen, salt bridges, van der Waals, pi stacking as well as hydrophobic events. Fig. 5 displays high level statistics as well as individual examples drawn from the distribution. Because proteins are not static molecules it is hard to place a threshold on how many interactions are required between domains, especially for samples generated unconditionally. Are a rule of thumb for setting a threshold we identified PDB ID: 3DH3 which contains two functional domains: binding and catalytic [1], while getcontacts identifies just under 20 interactions. Overall two domain BFN samples display good overall coherence with some examples even exhibiting discontinuous domains (**17693**, **16494**, **7980**, **437**). A more in-depth structural analysis is beyond the scope of this research article.

#### D. AbBFN generates natural heavy chains

Similarly to ProtBFN, an important goal of AbBFN is to learn the distribution of natural VH chains. We employ various measures of naturalness for unconditional heavy chains, considering the distribution of sequence lengths, amino acids, ESM plDDT. To this end, Fig. 6 presents a selection of such measures for 10,000 generated sequences in comparison to 10,000 natural sequences taken from OAS, from which we can infer that AbBFN closely matches the natural OAS distribution on which it was trained. As an additional measure, we align

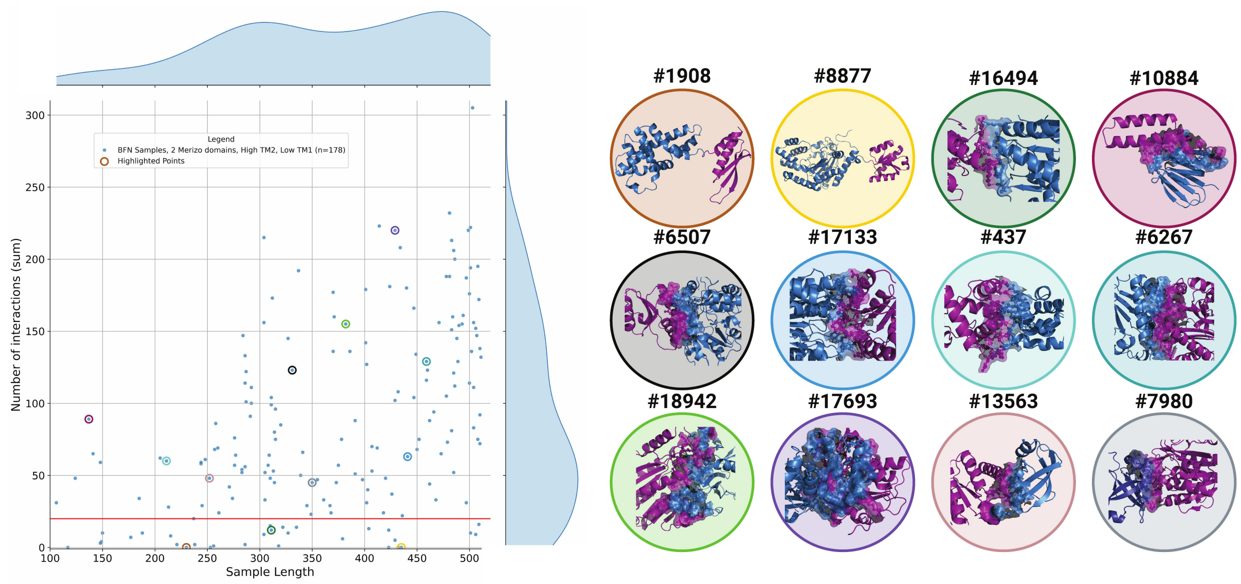


**Figure 5** | **Global coherence of two domain BFN samples.** Joint plot of BFN samples length vs Number of Interactions. The interactions considered are: hydrogen, salt bridges, van der Waals, pi as well as hydrophobic events. Highlighted points are displayed in. Each domain is coloured differently, and the overall structure is presented in a cartoon representation. The interface is presented using a surface representations while each individual interface residues is displayed using the ball and stick representation. Black dashed lines represent all the interactions identified by PyMOL [3] between the domains. Note that interactions at domain boundaries have been excluded.

generated structures, as predicted by ESMFold, against reference chain 6vi2_chain_B using TM-Align [9] to obtain a TM-score. We again observe strong concordance between AbBFN generated chains and natural OAS chains, and consistently high TM scores indicating that the generated sequences structurally resemble a canonical VH fold.

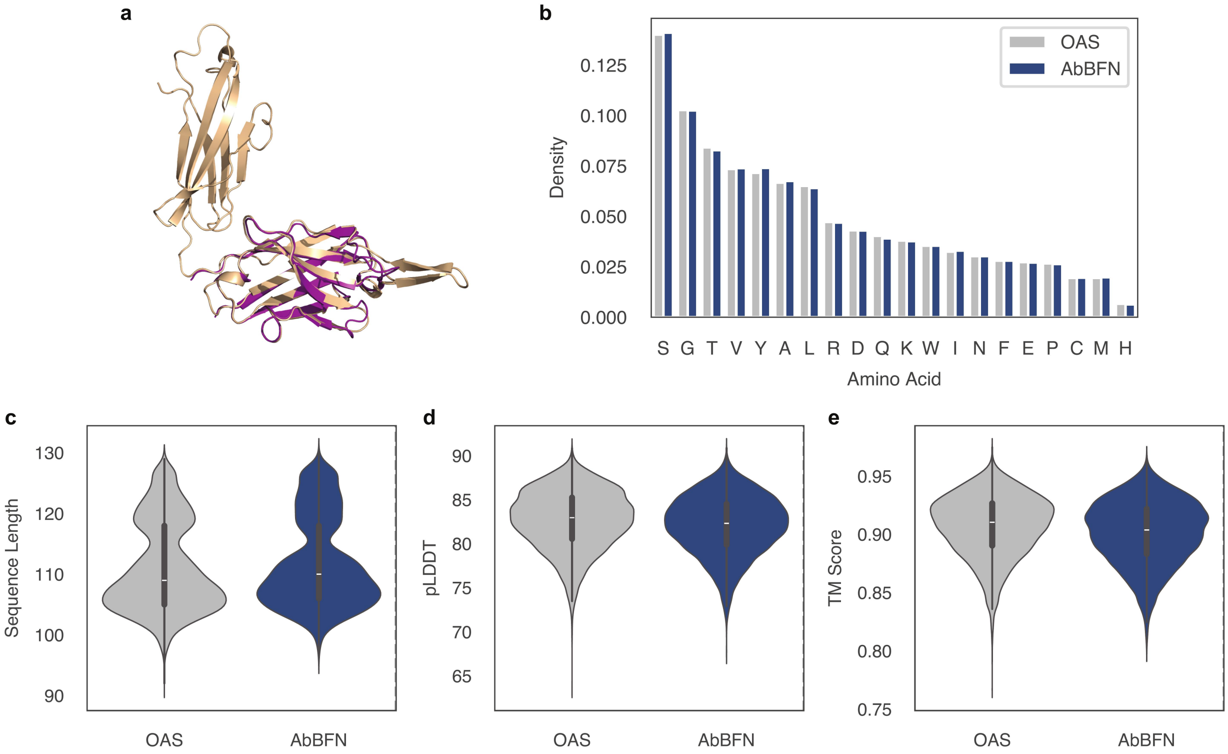


**Figure 6 | AbBFN is an effective *de novo* generator of VH chain sequences; generating novel VH chain sequences that in distribution under both local and global metrics.** In (**a**) a generated VH chain is compared to reference chain 6vi2_chain_b, showing clear structural consistency. The amino acid (**b**) frequencies show a strong match to the training distribution of the model. Both the sequence lengths (**c**) and predicted mean local distance difference test (pLDDT) scores from ESMFold [7] (**d**) are shown. Predictions with pLDDT>70 are generally considered high confidence. By computing the TM-score of generated and natural VH chains, against reference chain 6vi2_chain_b, we again observe strong consistency between AbBFN and OAS (**e**). In all cases, AbBFN is seen to well match the natural training distribution OAS.

[8] G. Porwal, S. Jain, S. D. Babu, D. Singh, H. Nanavati, and S. Noronha. Protein structure prediction aided by geometrical and probabilistic constraints. *Journal of Computational Chemistry*, 28(12):1943–1952, 2007.

[9] Y. Zhang and J. Skolnick. Tm-align: a protein structure alignment algorithm based on the tm-score. *Nucleic acids research*, 33(7):2302–2309, 2005.
